# Refining the pneumococcal competence regulon by RNA-sequencing

**DOI:** 10.1101/497099

**Authors:** Jelle Slager, Rieza Aprianto, Jan-Willem Veening

## Abstract

Competence for genetic transformation allows the opportunistic human pathogen *Streptococcus pneumoniae* to take up exogenous DNA for incorporation into its own genome. This ability may account for the extraordinary genomic plasticity of this bacterium, leading to antigenic variation, vaccine escape, and the spread of antibiotic resistance markers. The competence system has been thoroughly studied and its regulation is well-understood. Additionally, over the last decade, several stress factors have been shown to trigger the competent state, leading to the activation of several stress response regulons. The arrival of next-generation sequencing techniques allowed us to update the competence regulon, the latest report of which still depended on DNA microarray technology. Enabled by the availability of an up-to-date genome annotation, including transcript boundaries, we assayed time-dependent expression of all annotated features in response to competence induction, were able to identify the affected promoters and produced a more complete overview of the various regulons activated during competence. We show that 4% of all annotated genes are under direct control of competence regulators ComE and ComX, while the expression of a total of up to 17% of all genes is, either directly or indirectly, affected. Among the affected genes are various small RNAs with an as-of-yet unknown function. Besides the ComE and ComX regulons, we were also able to refine the CiaR, VraR (LiaR) and BlpR regulons, underlining the strength of combining RNA-seq with a well-annotated genome.

## INTRODUCTION

*Streptococcus pneumoniae* (the pneumococcus) is a mostly harmless human commensal found in the nasopharynx. However, when the pneumococcus leaves the nasopharynx and ends up in other niches, it may cause severe diseases, such as sepsis, pneumonia and meningitis (1). Especially among individuals with a weakened immune system, these diseases lead to over a million deaths per year (2). Although both vaccination and antibiotic therapy have been used successfully for, respectively, prevention and treatment of infections, the pneumococcus remains a threat to human health. This persistence is largely due to the remarkable genomic plasticity of the pneumococcus, allowing the acquisition of antibiotic resistance and evasion of the host immune response. Horizontal gene transfer, underlying the vast majority of such diversification strategies, is facilitated by pneumococcal competence. The competent state allows cells to take up exogenous DNA and integrate it into their own genome (i.e. transformation). During competence, various functionalities are activated, including DNA repair, bacteriocin production and several stress-response regulons (3, 4). This diversity of activated functions is relevant in light of the fact that a broad spectrum of antimicrobial compounds (causing various forms of stress) can actually induce competence development (5–7), through at least three distinct mechanisms: HtrA substrate competition (8, 9), *oriC*-proximal gene dosage increase (6) and chaining-mediated autocrine-like signaling (7). Other parameters that affect competence development include pH, oxygen, phosphate and diffusibility of the growth medium (10–12). The fact that various forms of stress induce competence, including several stress-response regulons, has led to the hypothesis that competence in the pneumococcus may function as a general stress response mechanism (13, 14).

Among the genes activated during competence are the CiaR and VraR (LiaR) regulons. Although the underlying molecular mechanisms of activation are unknown, both regulons have been associated with cell wall damage control (3, 15). Indeed, a growth lag during competence (4) and the reduced fitness of both *ciaR* and *vraR* mutants (3, 15) indicate that competence represents a significant burden for a pneumococcal cell. It seems plausible that the production and insertion of the DNA-uptake machinery (16) into the rigid cell wall has a significant impact on cell wall integrity. The CiaR regulon seems to be responsible for resolving such issues and preventing subsequent lysis (3). An additional dose of competence-related cell wall stress is caused by fratricide, where competent cells kill and lyse non-competent sister cells and members of closely related species. Specifically during competence, pneumococci produce a bacteriocin, CbpD, and the corresponding immunity protein, ComM (17). Secreted CbpD, aided by the action of autolysins LytA and LytC, can kill non-competent, neighboring cells, which then release their DNA and other potentially valuable resources. Eldholm et al. showed that the VraR regulon represents a second layer of protection, on top of ComM, by which competent cells prevent CbpD-mediated lysis (15). ComM is also crucial in causing cell division arrest during competence by inhibiting initiation of division and by interfering with the activity of StkP (18).

The regulation of competence (**Figure 1**) depends on the action of two key transcriptional regulators, ComE and ComX. The competence regulon is divided into early (i.e. ComE-dependent) and late competence (i.e. ComX-dependent) genes. Specifically, early competence involves, among others, the *comCDE* and *comAB* operons. A basal expression level of *comCDE* (19) ensures the production of the small peptide ComC, which contains a double-glycine leader and is processed and exported into the extracellular milieu by the bipartite transporter ComAB (20, 21). The resulting 17-residue matured peptide is referred to as competence-stimulating peptide (CSP) (22) and can interact with ComD, the membrane histidine kinase component of the two-component system ComDE (23). Upon CSP binding, ComD autophosphorylates and, subsequently, transfers its phosphate group to its cognate response regulator ComE (24). Finally, phosphorylated ComE dimerizes and binds specific recognition sequences to activate the members of the early-*com* regulon (25, 26). This regulon contains both the aforementioned *comAB* and *comCDE* operons, creating a positive-feedback loop that self-amplifies once a certain threshold of extracellular CSP is reached. Additional members of the early-*com* regulon are *comX1* and *comX2*, two identical genes that encode the alternative sigma factor ComX (σ^X^) (27). The rapid accumulation of ComX during early competence leads to the activation of promoters with a ComX-binding motif, resulting in the expression of the late-*com* regulon (3, 4, 28, 29). While the backbone of this regulatory system is quite well-understood, there are many other factors that complicate the matter, including the system’s sensitivity to growth medium acidity, potential repression of early-*com* genes by unphosphorylated ComE and DprA (26) and potential sRNA-mediated control of ComC expression (30). Finally, within 20 minutes after the initiation of competence, the process is largely shut down through a combination of different mechanisms (31–33).

**Figure 1.**
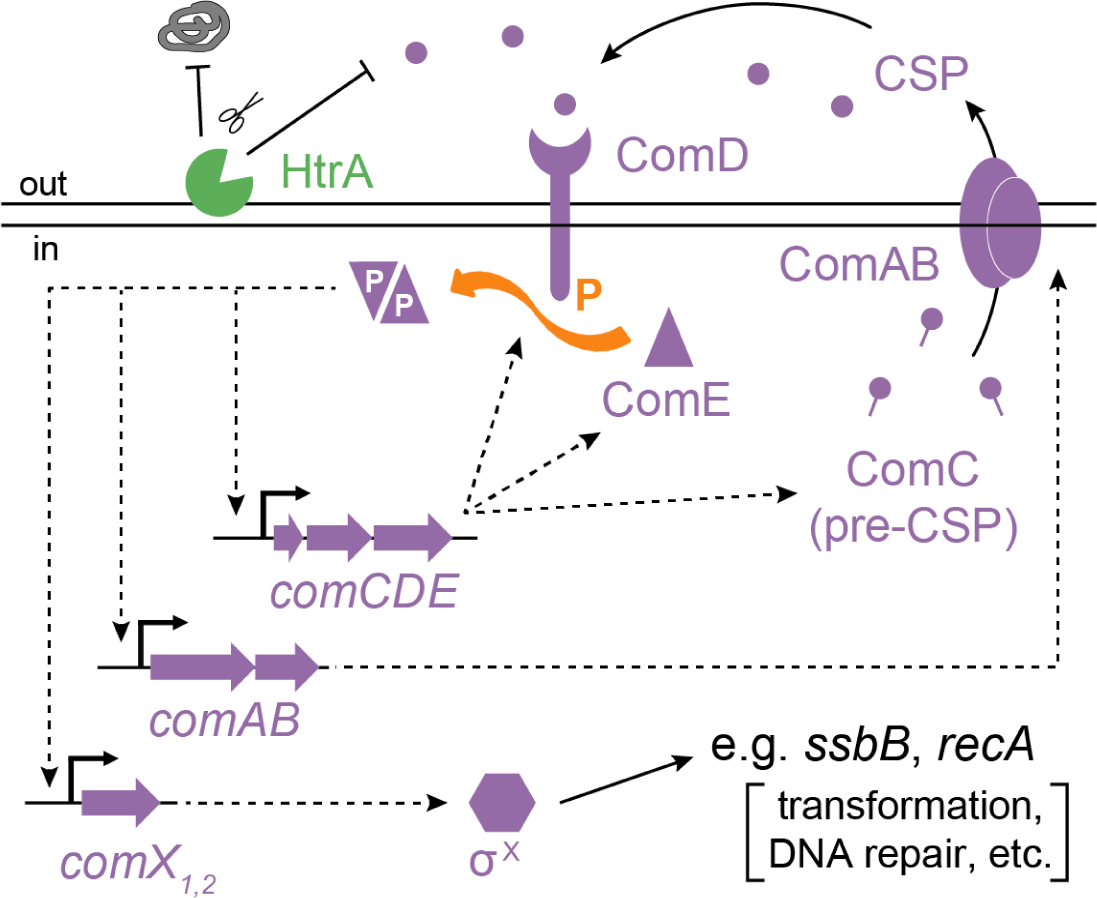
Overview of the regulatory network driving competence in *Streptococcus pneumoniae*. Adapted from Slager et al., 2014 (6).

To fully understand the implications of competence activation in the pneumococcus, it is important to know which genes are, directly or indirectly, differentially expressed during competence. Several comprehensive studies, based on DNA microarray technology, have been performed to determine the competence regulon, resulting in more than 100 reported competence-associated genes (3, 4, 34). All of these studies showed a high level of agreement on a certain core regulon, but discrepancies remained. Moreover, the recent identification of early-competence protein BriC (35, 36) illustrates that the description of the competence regulon can still be refined. In order to generate a completer and more nuanced overview of the competence regulon, we utilized data from PneumoExpress (35), a resource containing data on the pneumococcal transcriptome in various infection-relevant conditions. We used RNA-seq data sets from *S. pneumoniae* D39V (37) cells just prior to (t=0) and 3, 10 and 20 minutes after the addition of exogenous CSP. More importantly, compared to previous genome-wide assays of the competence regulon, which were based on DNA microarrays, our data set has higher sensitivity and precision and has a larger dynamic range (38). Secondly, the recent reannotation of the pneumococcal genome has revealed previously non-annotated protein-encoding sequences and small RNAs (37). DNA microarray studies are limited to the target sequences present on the array and a new data set was therefore required to obtain information on the expression of these new elements. Finally, the new annotation also contains information on transcription start sites (TSSs) and terminators (37), which allows both for a more accurate search of transcription regulatory motifs (e.g. ComE-or ComX-binding sites) and for the integration of operon information into the interpretation of transcriptome data.

As expected, our results largely confirmed previous microarray-based studies and we observed distinct time-dependent expression patterns of ComE-and ComX-regulated genes. In addition, we provide an overview of the transcription start sites most likely to be responsible for the observed transcriptome changes, adding up to, among others, 15 ComE-regulated, 19 ComX-regulated, 18 CiaR-regulated and 4 VraR-regulated operons. We identified 7 new non-coding RNAs, affected by several regulators, among the differentially expressed genes, but their role in competence requires future studies.

## MATERIALS AND METHODS

Here-studied samples are a subset of the data set presented in PneumoExpress (samples *C+Y*; *CSP, 3 min*; *CSP, 10 min*; *CSP, 20 min*; (35)) and detailed procedures regarding bacterial growth, RNA isolation, sequencing and read mapping are reported therein. The key points of these methods are summarized below.

### Culturing and harvesting of *S. pneumoniae* D39V

Eight tubes with 2 mL C+Y medium (pH 6.8, non-permissive for natural competence; (35)) without antibiotics were each inoculated with wild-type *S. pneumoniae* D39V cells (initial OD_600nm_ ∼ 0.004). When the cultures reached an OD_600nm_ of 0.05, two cultures were harvested for RNA isolation (t=0). To the other six, 100 ng/mL synthetic competence-stimulating peptide (CSP-1), purchased from GenScript (Piscataway, NJ), was added. Duplicate samples were harvested 3, 10 and 20 minutes after CSP-1 addition. Before harvesting, cultures were pre-treated with a saturated ammonium sulfate solution (39) to prevent protein-dependent RNA production and degradation. Afterwards, cells were harvested by centrifugation (20 min, 4 °C, 10,000 × g) and cell pellets were snap-frozen with liquid nitrogen and stored at −80 °C.

### Total RNA isolation, library preparation, sequencing and read mapping

RNA was isolated using phenol-chloroform extraction, followed by DNase treatment and another round of phenol-chloroform extraction (35). The quantity and quality of total RNA were estimated by Nanodrop, while a 1% bleach gel (40) was employed to confirm the presence of rRNA bands (23S, 2.9 kbp and 16S, 1.5 kbp) and absence of genomic DNA. Subsequently, RNA quality was again checked using chip-based capillary electrophoresis (Agilent Bioanalyzer). Stranded cDNA library preparation was performed, without depletion of ribosomal RNA, using the TruSeq^®^ Stranded Total RNA Sample Preparation Kit (Illumina, US). Sequencing was performed on an Illumina NextSeq 500, in 75 nucleotide single-end mode. The raw FASTQ data are accessible at http://www.ncbi.nlm.nih.gov/geo/ with accession number GSE108031 (samples B05-B11).

After a quality check with FastQC v0.11.5 (41), reads were trimmed using Trimmomatic 0.36 (42). Alignment of trimmed reads to the reference *S. pneumoniae* D39V genome (GenBank CP027540; (37)) was performed with STAR (43).

### Read quantification and differential gene analysis

The aligned reads were then counted (44) according to the D39V annotation file (GenBank CP027540; (37)) in a strand-specific fashion, allowing mapping to multiple sites (-M), for which fractional counts are reported (--fraction), and allowing reads to overlap multiple features (-O) to account for polycistronic operons.

Subsequently, we analyzed the libraries in R-studio (R v3.4.2). We performed differential gene expression analysis on rounded raw count by DESeq2 (45). Normalized expression levels are presented as TPM (transcripts per million) (46) and can be found in **Table S1**. Genes with a more than twofold absolute change of expression and a corresponding *p*-value of below 0.001 were considered to be significantly differentially expressed.

When possible, PneumoBrowse (https://veeninglab.com/pneumobrowse; (37)) was used to trace back differential expression of individual genes to a specific TSS and promoter region. As a starting point, the operon prediction from PneumoBrowse was used to define groups of genes differentially expressed in competence. It is important to note that strong transcriptional responses such as those observed during competence may have significant downstream effects. Even in the presence of highly efficient transcriptional terminators, which were defined to be operon boundaries in PneumoBrowse, such read-through effects may be visible. Therefore, these co-expressed groups were refined by inspection of the raw data in PneumoBrowse and the consideration that minor read-through from a highly expressed gene can still be significant if the expression of the downstream gene is sufficiently lower.

### Clustering and creation of position weight matrices

Using the weighted gene co-expression network analysis (WGCNA) R software package (47), genes were clustered based on their *rlog* (regularized log) expression value (**Table S2**), as output by DESeq2, across all 22 infection-relevant conditions analyzed in PneumoExpress (35). We noticed that the reported members of the ComE (25, 26), ComX (3, 4, 34) and CiaR (30, 48, 49) regulons each largely ended up in specific clusters (here: clusters 29, 11 and 33 for ComE, ComX and CiaR, respectively). Reported regulon members that properly clustered in these three identified main clusters, which will be referred to as ‘training sets’ (**Table S3**), were used to define the recognition motifs of these three regulators, in the form of position weight matrices (PWMs), and to determine the optimal distance of such a motif from the TSS. Using the MEME suite (50), we analyzed the upstream regions of each training set for enriched sequence motifs. Firstly, since earlier work showed slightly different consensus sequences for the two tandem ComE-boxes that make up the ComE-site (26), we extracted the left ComE motif (CEM_L_) by scanning the regions from 77 to 63 bps upstream and the right motif (CEM_R_) by scanning the regions 56 to 42 bps upstream of TSSs in the training set. The ComX-binding motif (CXM) was determined from the regions 35 bps upstream to the +1 site (TSS). When building CEM_L_, CEM_R_ and CXM PWMs, each sequence in the training set was required to have exactly one match to the motif, in the transcription direction (i.e. on the locally defined ‘plus’-strand). CiaR has been described to bind to a direct repeat (30) and we scanned the regions 41 to 19 bps upstream, allowing for multiple hits per sequence in the training set. While some members of the CiaR regulon have binding sites on the opposite strand, none of these genes were part of the training set and the CiaR-binding motif (CRM) were therefore also limited to the ‘plus’-strand. Genes reported to belong to the VraR (LiaR) regulon (15) did not cluster together throughout the 22 conditions and for some of these genes the TSS was unknown. To be able to extract a VraR-binding motif (VRM), we combined upstream regions of the pneumococcal genes *spxA2* (SPV_0178), *vraT* (SPV_0350) and SPV_0803 with those of six *Lactococcus lactis* genes that were reported to be regulated by close VraR homolog CesR (15, 51): llmg_0165, llmg_0169, llmg_1115, llmg_1155, llmg_1650 and llmg_2164. Cappable-seq (52) was used to identify *L. lactis* TSSs (S.B. van der Meulen and O.P. Kuipers, unpublished). Importantly, we did not use the standard ‘0-order model of sequences’ as a background model for motif discovery, but instead created background models corresponding to the corresponding regions upstream of all known TSSs in the pneumococcal genome (e.g. −35 to +1 for ComX). Additionally, we defined summary consensus sequences using IUPAC nucleotide coding. Since the CiaR-binding motif reportedly consists of two perfect repeats, we determined the consensus based on the 16 motif occurrences in the CiaR training set (8 promoter sequences). Single base codes (A, C, G, T) were called when 75% (rounded up) of all promoters matched. Double base codes (R, Y, S, W, K, M) were called when 8/9 (ComE and VraR), 15/16 (ComX), 5/5 (BlpR) promoters matched either of the two encoded bases. Triple base codes (B, D, H, V) were called when all promoters matched either of the three encoded bases. Note that, due to its degenerate appearance, the *blpRS* promoter was excluded when determining the BlpR-binding consensus.

### Assigning putative regulons

After creating PWSs for ComE-, ComX-, CiaR-and VraR-binding sites, we used FIMO (53) to scan the 100 bps upstream of all known pneumococcal TSSs for matches to these motifs. Here, too, we used the appropriate background models (see above). A cutoff *q*-value of 0.01 was used for hits with ComX-and VraR-binding motifs. We defined a reliable ComE-binding site as CEM_L_-[N_11-13_]-CEM_R_, using a cutoff *p*-value of 0.01 for each motif. Similarly, we defined a CiaR-binding site as CRM-[N_5-6_]-CRM. Additionally, to assign a gene cluster to a certain putative regulon, we also put a constraint on the position of the motif relative to the corresponding TSS, based on the typical spacing observed in the training sets. Thus, the allowed first nucleotide positions were [-77/-76/-75/-74/-73] for ComE, [-30/-29/-28] for ComX, [-40/-39/-38/-37/-36] for CiaR, and [-51/-50/-49/-48/-47] for VraR.

Putative binding sites for other regulatory proteins were copied from the propagated *S. pneumoniae* D39 regulons, as found in the RegPrecise database (54) and annotated in PneumoBrowse (37). RNA switches, annotated in D39V, were also taken into consideration as putatively responsible regulatory mechanisms.

### Gene enrichment analysis

Differentially expressed genes that could not be ascribed to the action of ComE, ComX, CiaR or VraR, were subjected to gene enrichment analysis (functional analysis). For this, Gene Ontology and KEGG classifications were extracted from the GenBank file corresponding to the latest annotation of *S. pneumoniae* D39V (37). Additionally, predicted transcription factor binding sites were used to assign genes to their putative regulons. A total of 448 hypergeometric tests were performed and a Bonferroni-corrected cutoff *p*-value of 0.0001 (i.e. 0.05 divided by 448) was used to determine whether certain regulons or Gene Ontology or KEGG classes were overrepresented among differentially expressed genes. We excluded overrepresented classes when all affected genes belonged to the same operon, since the activation of a single promoter does not confer any statistical evidence.

## RESULTS

### Competence induction disrupts the pneumococcal transcriptional landscape

Differential gene expression analysis revealed that many genes (13-17%; from gene-based or promoter-based analysis, respectively) are affected by the induction of competence (**Figure 2**): a total of 288 genes undergo a change in expression of more than twofold. Out of these, 192 genes are exclusively upregulated, 94 are exclusively downregulated and 2 genes are upregulated at one time point and downregulated at another. When using a stricter fold change cutoff of fourfold, 141 genes are still significantly affected, 119 of which are up-and 22 are downregulated. As can be seen in **Figure 2**, upregulated genes tend to be affected more strongly and consistently, while not a single gene is significantly downregulated in all three time points.

**Figure 2.**
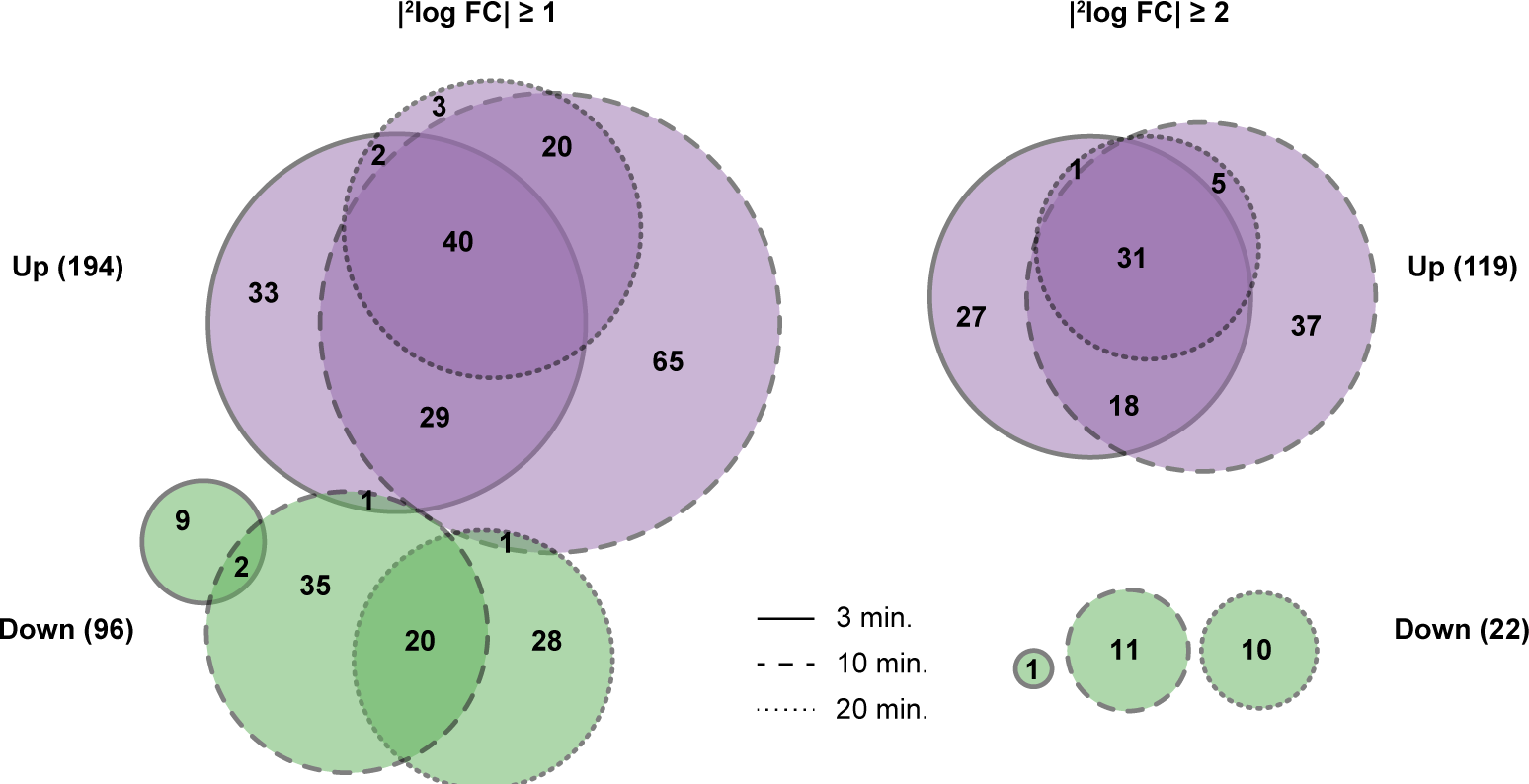
Venn diagrams of differentially expressed genes, created with http://eulerr.co. Diagrams show how many genes were significantly up-(purple) or downregulated (green), using a cutoff fold change of 2 (left) or 4 (right). Differential expression 3, 10 and 20 minutes after addition of CSP is indicated by solid, dashed and dotted lines, respectively.

### Identification of ComE-, ComX-and CiaR/VraR-regulated WGCNA clusters

WGCNA clustering (see **Materials and Methods**) of all genes, based on their regularized log (rlog) expression levels across the 22 conditions included in PneumoExpress (35), yielded 36 clusters (**Table S2**). Using these results, we verified whether some of the clusters corresponded to specific regulons known to be affected during competence. Indeed, one of these clusters (cl. 29, n=26) contained 20 out of 25 genes that have been previously reported to be regulated by ComE (3, 4, 34), including *briC* (SPV_0391), which was only recently identified as a member of the competence regulon (35, 36). The five ComE-regulated genes that did not end up in this cluster include *blpA, blpY, blpZ* and *pncP* (SPV_0472-75), which are part of the BlpR regulon and whose promoters are likely to have lower affinity for ComE (55, 56). The fifth off-cluster ComE-regulated gene is *ybbK* (SPV_1984). A second cluster (cl. 11, n=56) contained 41 out of 51 members of the reported ComX regulon (3, 4, 34), confirming the power of the WGCNA approach, while simultaneously highlighting the general reliability of previous descriptions of the competence regulon. Less clearly, 13 out of 32 known CiaR-regulated genes (30, 48, 49) and 5 out of 14 VraR-regulated genes (15) clustered together (cl. 33, n=22). The fact that genes from the CiaR and VraR regulon cluster less clearly may, in part, be explained by the more diverse nature of their regulation. For example, the heat-shock *hrcA*-*grpE* operon is not only regulated by VraR, but also by HrcA itself, accounting for a different expression dynamic across the diverse conditions sampled for PneumoExpress (35). Additionally, the two TSSs of *tarIJ*-*licABC* (SPV_1127-23) (30) and the downregulatory effect of CiaR on the *manLMN* operon (SPV_0264-62) (49) may prevent clear clustering.

### Time-resolved expression profiles of several regulons during competence

We visualized the typical time-resolved expression patterns of the various regulons, where we plotted the fold changes, relative to t=0, of all genes that were i) previously reported to be activated by the corresponding regulator, and ii) fell into the associated WGCNA cluster (see above). We refer to these sets of genes as ‘core members’ of their respective regulons. It is clear from these plots that ComE-regulated genes peak early and rapidly drop in expression level afterwards (**Figure 3A**, left). This is in line with previous studies, which showed that early competence is actively shut down through the action of late-competence protein DprA. By specifically binding to active, phosphorylated ComE, DprA causes a shift towards a state where regulated promoters are, instead, bound by dephosphorylated ComE, leading to a shutdown of transcription (33, 57).

**Figure 3.**
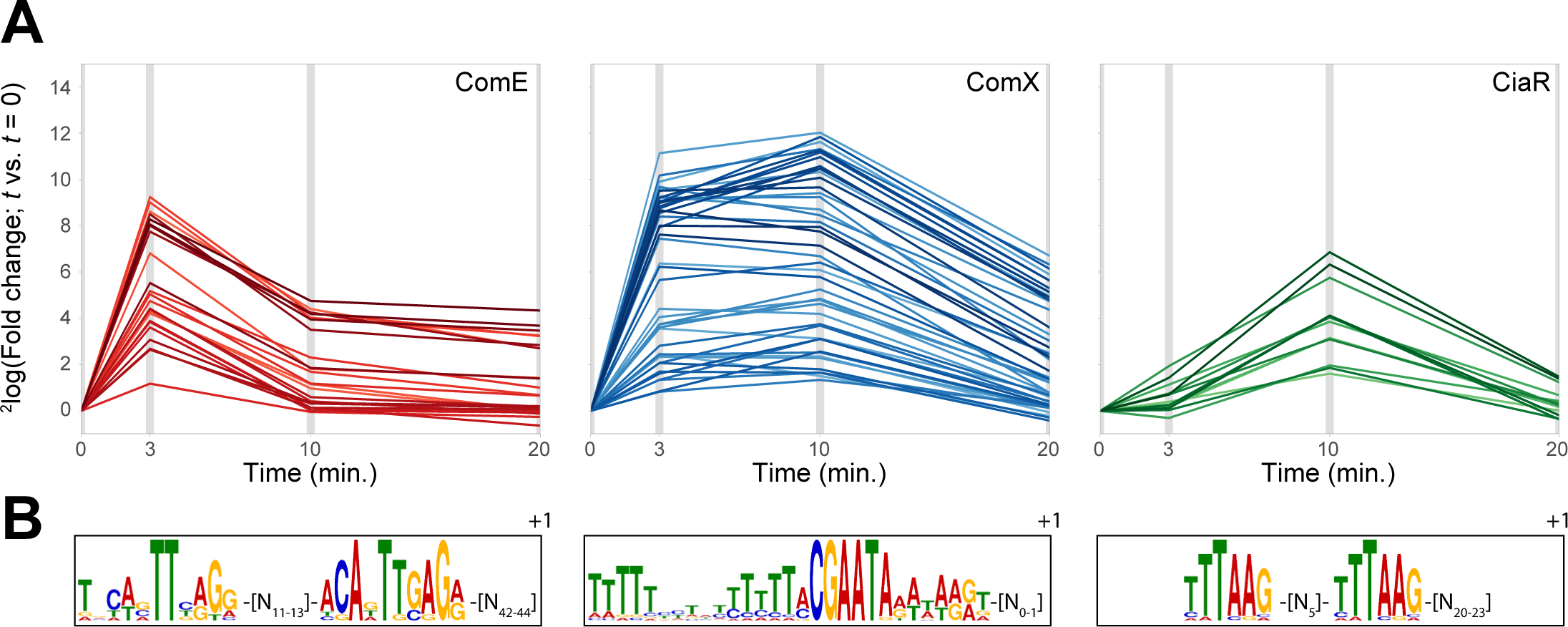
Affected regulons during competence. (**A**) Expression profiles (fold-changes vs. t=0) of genes previously reported to be ComE-, ComX-or CiaR-activated. Only genes that fell into the appropriate WGCNA cluster (cl. 29 for ComE, cl. 11 for ComX, cl. 33 for CiaR) were included. (**B**) Position weight matrices of the recognition sites for ComE, ComX and CiaR, as determined with MEME (50), from the promoters of the core members of the corresponding regulons.

Similar to the ComE regulon, the expression of ComX-regulated genes also increases rapidly, with high fold changes after 3 min. However, unlike the ComE regulon, the expression of these genes remains stable for a longer period of time, with most still increasing their level until 10 min. after CSP addition (**Figure 3A**, center). The following decrease in expression level is, in part, indirectly linked to the shutdown of early competence, since both production and stabilization by ComW (31) of ComX depend on the activity of phosphorylated ComE. However, a recent modelling approach suggested that another shutdown mechanism was required to explain the observed rate of late competence shutdown (33). The authors argue, convincingly, that competition between sigma factors ComX (σ^X^) and RpoD (σ^A^) for interaction with RNA polymerase and/or stabilizing factor ComW would be a suitable explanation for the discrepancies between the model and experimental data. Indeed, the fact that *rpoD* is upregulated up to tenfold during competence (3, 4) would make this a credible hypothesis, although *rpoD* upregulation could also simply serve to restore the expression levels of RpoD-controlled genes.

Constituting a more indirect consequence of competence induction, the CiaR-mediated response is generally weaker and delayed, compared to the ComE and ComX regulons (**Figure 3A**, right). Interestingly, the activation of this regulon also seems to be quite transient of nature, with a fast drop in expression from 10 to 20 min. after CSP addition.

Finally, the expression profile of all reported VraR-regulated genes (15), regardless of their clustering behavior throughout infection-relevant conditions, was similar to that of the CiaR regulon (**Figure 4A**).

**Figure 4.**
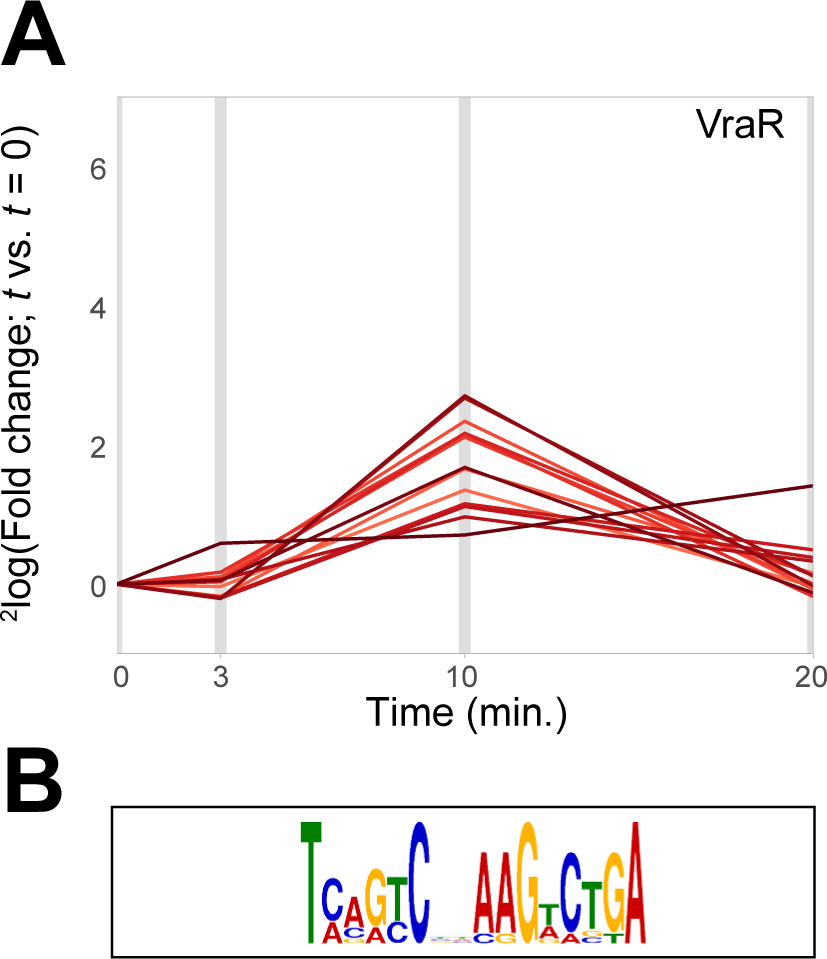
VraR (LiaR) regulon. **A**. Expression profiles (fold-changes vs. t=0) of genes previously reported to be VraR-activated. **B**. Position weight matrix of the recognition site for VraR, as determined with MEME (50), from the promoters of three pneumococcal VraR-regulated operons and 6 *L. lactis* promoters (15).

### Shared expression trends within operons allow switching from gene to promoter level analysis of transcriptional regulation

Transcriptome studies are typically performed on a per-gene level, reporting for each individual gene whether or not it is differentially expressed between the studied conditions. That type of information is certainly relevant when trying to assess what changes occur in a cell or population when confronted with a certain change in environment or identity. However, to find out *how* these changes have come about, it may also be interesting to consider which transcripts or, rather, which promoters have been affected. Therefore, we will, where possible, use our previously created map of the pneumococcal transcriptional landscape (35, 37) to identify which promoters are responsible for the observed differential expression of individual genes. As an example, we highlight a cluster of 22 genes (SPV_0192-213), encoding 21 ribosomal proteins and protein translocase subunit SecY (**Table 1**). Although the entire operon was reported to be downregulated by Peterson et al. (4), we only observed significant upregulation for 10 genes (just above the 2-fold cutoff) and no significant effects for the rest of the operon. However, both from visual inspection and the fact that all genes cluster together in the WGCNA analysis, it is clear that the entire operon behaves as a single transcriptional unit, with a modest upregulation 3 minutes after CSP addition, followed by a drop in expression at 10 minutes. Therefore, regulation of a single TSS, at 195877 (+), would suffice to explain the behavior of these 22 genes.

**Table 1.**
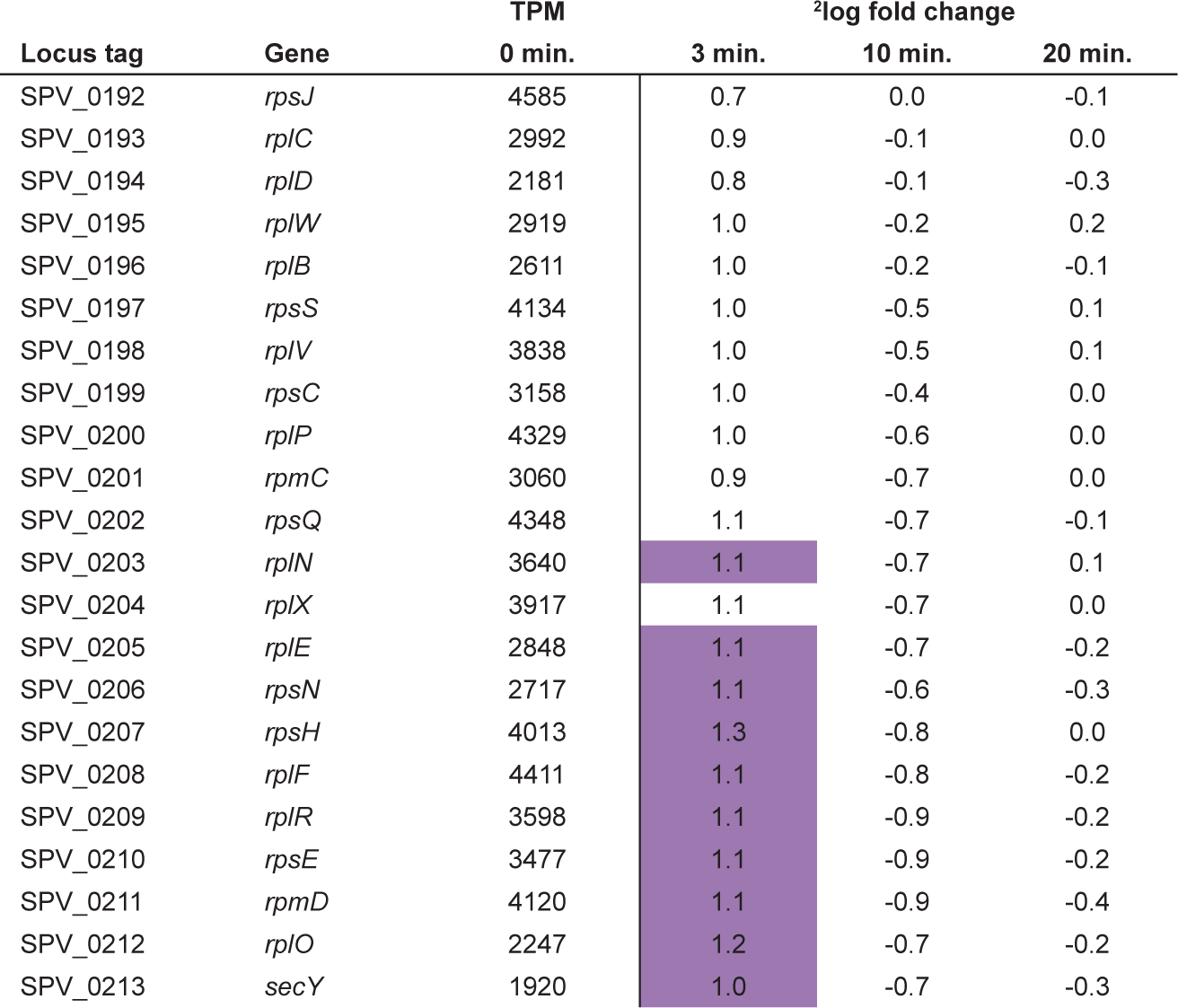
Expression trend of a 22-gene operon (SPV_0192-213) encoding 21 ribosomal proteins and protein translocase subunit SecY, transcribed from TSS 195877 (+). Although not all genes pass the significance cutoffs, they clearly share the same time-dependent expression pattern. Purple cells indicate significance (*p* < 0.001, ^2^log FC > 1; DESeq (45)).

### Early competence genes: the ComE and BlpR regulons

We reasoned that the promoter regions of core members (defined above) of a regulon were likely to yield a more reliable consensus binding site of the corresponding regulator, compared to when such a consensus were based on all known regulated genes, as is the common procedure. Therefore, combining transcription start site (TSS) data on these selected genes (**Table S3**) as reported previously (37) and known characteristics of regulatory sites (e.g. typical distance from TSS), we redefined the binding motifs for ComE, ComX and CiaR (See **Materials and Methods** for details). The identified ComE-binding sequence (**Figure 3B**, left; **Figure 5A**) strongly resembles previous reports (25, 26) and consists of two imperfect repeats. The spacing between these repeats is 11-13 nts (mode=11), while the spacing between the right motif and the TSS is 42-44 nts (mode=42). Clearly, the second repeat is most conserved and is likely to be most important for ComE recognition. In summary, this yields the following consensus ComE-binding motif: [TNYWVTTBRGR]-[N_11_]-[ACADTTGAGR]-[N_42_]-[TSS] (**Figure 5A**).

**Figure 5.**
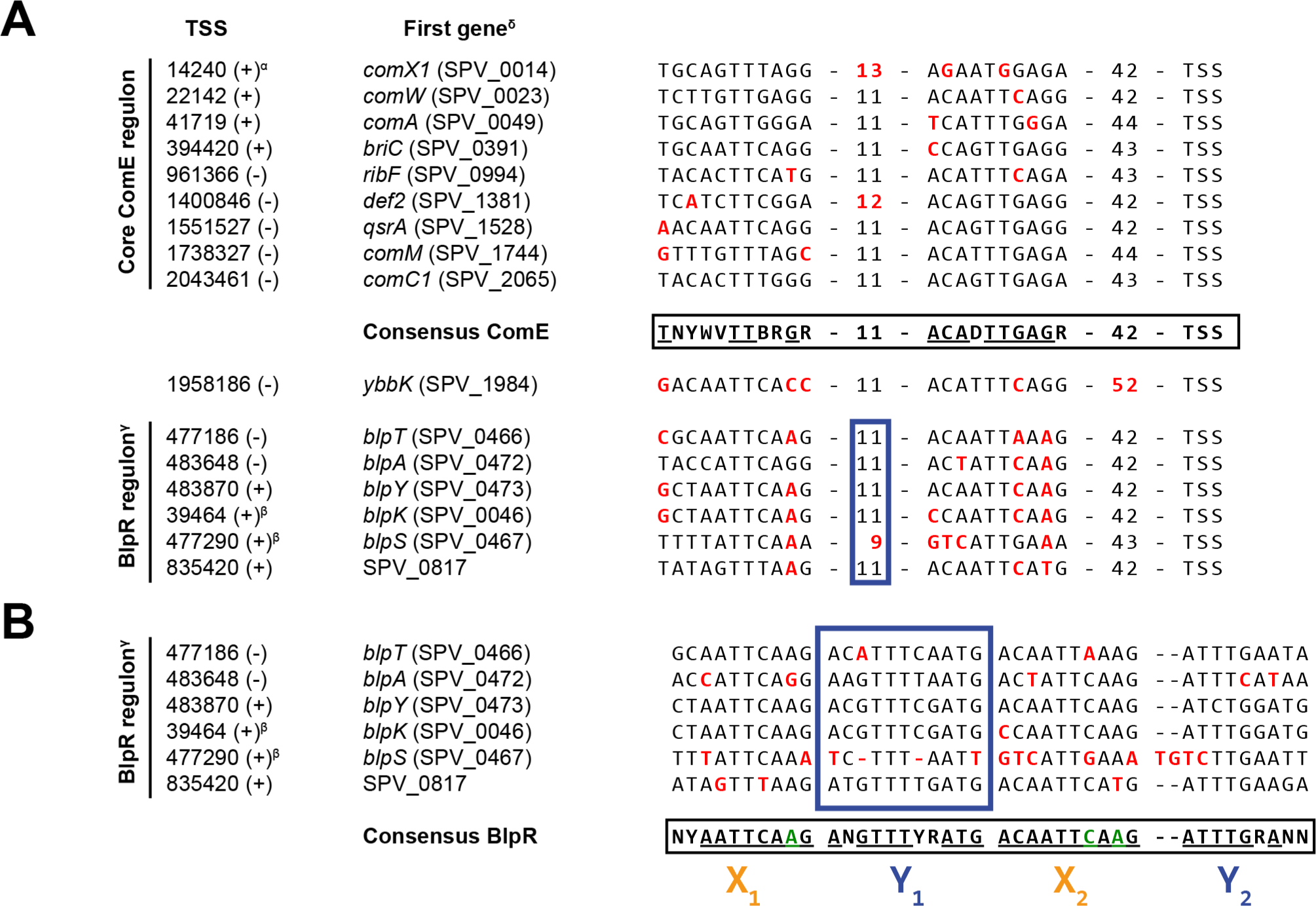
(**A**) ComE-binding sequences on the *S. pneumoniae* D39V genome. Consensus sequence was determined as described in **Materials and Methods**. Multiple possible nucleotides are indicated, according to IUPAC nomenclature, by R (A/G), Y (C/T), W (A/T), B (C/G/T), D (A/G/T), V (A/C/G) or N (any). Nucleotides and spacings colored red deviate from this consensus. (**B**) Putative BlpR-binding site consensus. The blue box corresponds to the internal spacer indicated in panel A. Consensus sequence (IUPAC nomenclature, as in **A**) was determined as described in **Materials and Methods**, where letters colored green indicate where the BlpR consensus is incompatible with the ComE consensus. ^α^Promoters of *comX1* (SPV_0014) and *comX2* (SPV_1818) are identical. ^β^These operons were not differentially expressed in response to CSP addition. ^γ^The genes encoding the export machinery of signaling peptide BlpC and bacteriocins BlpK and PncW, *blpA* and *blpB*, are frameshifted, eliminating the regulatory positive-feedback loop of the Blp system. ^δ^First gene both annotated in D39V and D39W (58).

As Martin et al. previously described (26), the internal spacing and the right arm of the binding sequence in P_*comX*_ deviate from the consensus (**Figure 5A**). Indeed, from our data, this deviation seems to lead to a somewhat lower expression level from P_*comX*_ (**Table 2**), which is partially compensated by the presence of two copies of *comX* on the chromosome. In contrast, we found no indication that mismatches with the consensus ComE-binding sequence found in the promoter region of *comAB* resulted in lower expression of those genes (**Table 2**), which was suggested by Martin et al. (26) to explain an earlier observation that *comAB* levels were rate-limiting in the development of competence (59). Indeed, although higher ComAB levels may indeed accelerate competence development, the fact that a duplication of *comC* (21, 24) leads to competence upregulation suggests that ComAB is not exporting CSP at maximum capacity in wild-type cells.

**Table 2.** ComE-regulated genes, distributed over 15 operons, as indicated by grey/white colored blocks. Members of the BlpR regulon are included, as indicated under ‘Notes’. Secondary (under ‘Notes’) indicates either read-through after incomplete termination or the influence of an additional TSS. For complete information, including TSS positions, see **Table S5**. Purple cells indicate significance (*p* < 0.001, ^2^log FC > 1; DESeq (45)). *Pseudogene.

| Gene | Product | TPM<br>0 min. | <sup>2</sup>log fold change |  |  | Notes |
| --- | --- | --- | --- | --- | --- | --- |
|  |  |  | 3 min. | 10 min. | 20 min. |  |
| *comX1* | Competence-specific sigma factor | 8 | 9.1 | 3.8 | 3.1 |  |
| *tRNA-Glu-1* | tRNA-Glu-UUC | 387 | 2.8 | 0.1 | -0.6 | Secondary |
| *comW* | Competence positive regulator | 17 | 9.1 | 4.7 | 3.0 |  |
| *purA* | Adenylosuccinate synthetase | 619 | 4.4 | 1.0 | 0.0 | Secondary |
| *ccnC* | csRNA3 | 155 | 4.4 | 1.2 | 0.0 | Secondary; also CiaR regulon |
| *srf-03* | ncRNA of unknown function | 39 | 7.1 | 2.0 | 1.5 |  |
| *comA* | CSP ABC transporter ATP-binding protein | 21 | 9.3 | 4.1 | 3.4 |  |
| *comB* | CSP ABC transporter permease | 24 | 9.5 | 4.4 | 3.3 |  |
| *briC* | Hypothetical protein | 97 | 4.9 | 1.7 | 0.7 |  |
| *ydiL\** | Putative membrane peptidase | 67 | 5.3 | 2.4 | 1.0 |  |
| *blpT* | BlpT protein | 8 | 3.9 | 0.7 | 1.5 | BlpR regulon |
| *blpC* | Peptide pheromone | 3 | 4.4 | 2.7 | 1.9 | BlpR regulon |
| *blpB\** | Peptide ABC transporter permease | 3 | 4.5 | 2.2 | 2.4 | BlpR regulon |
| *blpA\** | Peptide ABC transporter permease/<br>ATP-binding protein | 2 | 5.1 | 2.1 | 2.5 | BlpR regulon |
| *pncW* | Putative bacteriocin | 13 | 4.0 | 1.1 | 2.0 | BlpR regulon |
| *blpY* | Bacteriocin immunity protein | 17 | 4.3 | 1.4 | 2.2 | BlpR regulon |
| *blpZ* | Immunity protein | 13 | 4.3 | 1.5 | 1.9 | BlpR regulon |
| *pncP* | Putative protease | 14 | 4.2 | 1.3 | 1.9 | BlpR regulon |
| SPV_2249 | Hypothetical protein | 50 | 1.2 | 0.3 | 0.8 | BlpR regulon |
| SPV_0817 | CAAX amino terminal protease family protein | 45 | 1.3 | 0.2 | 0.4 | BlpR regulon |
| *ribF* | FMN adenylyltransferase/riboflavin kinase | 147 | 5.2 | 1.2 | 0.7 |  |
| *yaaA* | UPF0246 protein | 85 | 1.2 | 0.0 | -0.3 | Secondary |
| SPV_1379 | Hypothetical protein | 183 | 3.3 | 0.0 | -0.1 | Secondary |
| SPV_1380 | Cell shape-determining protein | 276 | 3.9 | 0.1 | 0.0 | Secondary |
| *def2* | Peptide deformylase | 177 | 3.7 | 0.0 | -0.1 |  |
| *srf-22* | ncRNA of unknown function | 54 | 4.1 | 0.2 | -0.7 | Secondary |
| *qsrB* | ABC transporter permease - Na<sup>+</sup> export | 216 | 4.5 | 0.4 | -0.2 | Secondary |
| *qsrA* | ABC transporter ATP-binding protein - Na<sup>+</sup> export | 223 | 4.5 | 0.6 | 0.2 | Secondary |
| *lytR* | Transcriptional regulator | 355 | 2.8 | 0.1 | 0.1 | Secondary |
| SPV_1742 | Acetyltransferase | 319 | 2.7 | 0.3 | 0.2 | Secondary |
| *tsaE* | tRNA processing protein | 280 | 3.1 | 0.3 | 0.2 | Secondary |
| *comM* | Immunity factor | 9 | 8.0 | 4.4 | 2.8 |  |
| *tRNA-Glu-3* | tRNA-Glu-UUC | 387 | 2.8 | 0.1 | -0.6 | Secondary |
| *comX2* | Competence-specific sigma factor | 8 | 9.1 | 3.8 | 3.1 |  |
| *ybbK* | Putative membrane protease subunit | 795 | 2.1 | 0.6 | 0.6 | TSS too far from ComE-binding site |
| *tRNA-Asn-2* | tRNA-Asn-GUU | 181 | 5.1 | 0.8 | 0.4 | Secondary |
| *tRNA-Glu-5* | tRNA-Glu-UUC | 63 | 5.8 | 1.9 | 1.5 | Secondary |
| *comE* | Two-component system response regulator | 38 | 8.3 | 4.0 | 3.5 |  |
| *comD* | Two-component system sensor histidine kinase | 27 | 8.4 | 4.3 | 3.8 |  |
| *comC1* | Competence-stimulating peptide precursor | 36 | 8.8 | 5.1 | 4.6 |  |

As reported before, the *comAB* genes are preceded by a BOX element (60), an imperfectly repeated DNA element occurring 127 times in the D39V genome (37). Interestingly, the BOX element is located downstream of the ComE-regulated TSS and therefore part of the *comAB* transcript. It was shown by Knutsen et al. that the BOX element upstream of *comAB* is important for the fine-tuning of competence (61), but the underlying mechanism is unknown. We previously detected several RNA fragments terminated between the BOX element and the start codon of *comA*, leading us to annotate it as a ncRNA, *srf-03* (37). Although some BOX elements were reported to contain putative protein-encoding sequences, we did not find any uninterrupted coding sequence in this specific case. It seems unlikely that the BOX element functions as an RNA switch, since *srf-03* and *comAB* displayed the same time-dependent expression patterns with very low expression at t=0 (**Table 2**). It is tempting to speculate that the prematurely terminated transcript, *srf-03*, plays a role in competence regulation. However, a similar effect on transcription was observed when the BOX element in front of *qsrAB* (also ComE-regulated) was removed (61) and we did not find any evidence for premature termination between that BOX element and the start of *qsrA*.

Analysis of all upregulated promoters resulted in the detection of five additional operons putatively regulated by ComE (the complete proposed ComE regulon is listed in **Table 2**), including *ybbK*, a known early-*com* gene. The weaker induction of this gene and the fact that its expression did not cluster with other early-*com* genes can be explained by the fact that the ComE site is located 10 nts too far from the TSS, compared to a canonical ComE-regulated gene (**Figure 5A**).

Three other ComE-induced operons (*blpT*; *blpABC*; *pncW*-*blpYZ*-*pncP*) are known to be part of the BlpR regulon and their activation is the result of crosstalk, where ComE can recognize the binding sites of BlpR, but with lower efficiency (55, 56). Indeed, close inspection of the corresponding promoter regions shows marked differences with those of other ComE-regulated genes, consistently deviating from the consensus ComE-binding site at specific positions (**Figure 5A**). The same discrepancies were observed in the promoter regions of *blpK* and, to a lesser extent, *blpSRH*, the two remaining *blp* operons. These operons were not differentially expressed during competence, probably due to the poorer resemblance to the ComE-binding consensus. Additionally, both *blpK* and *blpSRH* are constitutively expressed, such that any minor inducing effect by ComE would be negligible. A multiple sequence alignment of the five known *blp* operons in strain D39V revealed a conserved sequence very similar to, but slightly more extended than, the putative BlpR-binding site postulated by De Saizieu et al. (62). The here-reported binding site can be seen as an imperfect tandem 19-21 bp repeat: [NYAATTCAAGANGTTTYRATG]-[ACAATTCAAG(NN)ATTTGRANN]-[N_33_]-[TSS] More specifically, the region can be written as X_1_-Y_1_-X_2_-Y_2_, where X (resembling the ComE-binding site) and Y are 10 and 9-11 bps in length, respectively, having a highly conserved ‘TT’ (or ‘TTT’ in Y) at their centers (**Figure 5B**). Interestingly, the promoter region of the final operon putatively regulated by ComE, SPV_2249-SPV_0817, resembles the putative BlpR recognition site (**Figure 5B**) and we speculate that these genes are actually part of the BlpR regulon. This idea is supported by the very modest induction (2.3- and 2.5-fold, respectively) of this operon during competence. Additionally, SPV_0817 encodes a probable CAAX protease (37) that could be speculated to be involved in immunity against self-produced bacteriocins (63, 64).

Finally, only one feature from the WGCNA cluster associated with ComE regulation, remained that could not be directly linked to a ComE-binding site. This feature, a pseudogene (SPV_2414), is part of an ISSpn7 insertion sequence (65) and represents a truncated version of the gene encoding the corresponding transposase. Since the D39V genome contains eight additional sites with a ≥95% sequence identity, clearly undermining mapping fidelity, and no significant differential expression was observed in any competence time point, we ruled out SPV_2414 as a member of the ComE-regulon.

### Late competence genes: the ComX regulon

Directly following the strong, ComE-mediated increase in *comX* expression, the late-competence regulon is activated. Based on the promoter sequences of core members of the late-*com* regulon (**Table S3**, **Figure 6**), the ComX recognition sequence was re-evaluated. Not surprisingly, the identified motif (**Figure 3B**, center) does contain a near-perfect match to the previously reported 8-nucleotide consensus sequence (3, 28, 29): [TMCGAATA]. However, our analysis shows that the region relevant to ComX binding is likely much wider than that: with a thymine-rich stretch upstream and a, less-conserved, adenine-rich stretch downstream of the reported 8 nucleotides, the actual recognition site is extended to 20-30 basepairs. In summary, this yields the following consensus ComX-binding motif: [TTTTTNHNNNYTHT<u>TMCGAATA</u>DWNWRRD]-[TSS] (**Figure 6**). Besides the 16 core ComX-regulated operons, we identified three additional promoters containing the here-reported motif (**Table 3**). Firstly, SPV_0027-30, an operon encoding, among others, a dUTP pyrophosphatase (*dut*) and DNA repair protein RadA (*radA*), was already previously reported to be part of the late-*com* regulon (3, 4), but did not cluster with the core ComX regulon. A secondary TSS, 11 nucleotides downstream of the ComX-regulated TSS, could be responsible for the lower correlation with other ComX-regulated genes. Similarly, a second previously reported late-*com* gene, SPV_0683, may be under the control of a secondary, not yet identified TSS, besides the here-reported ComX-activated TSS (**Table 3**), as supported by its relatively high expression level prior to CSP addition and sequencing coverage observed in PneumoBrowse (37). A third ComX-binding site was found downstream of *prs1* (SPV_0033) and immediately upstream of a novel ncRNA, *srf-01* (SPV_2081), which we identified recently (37). While this addition to the known competence regulon seemed interesting at first, the partial overlap between the ncRNA and a nearby IS element (containing pseudogene SPV_0034) led us to question the functionality of this novel element (**Figure S1**). Additionally, another pseudogene (SPV_2082) was located on the other side of the IS element, also under the control of ComX (**Table 3**). A multiple genome alignment of several pneumococcal strains (not shown) revealed that the ComX-binding element downstream of *prsA1* (i.e. *prs1*), was conserved in, among other strains, *S. pneumoniae* INV200 (GenBank: FQ312029.1), but was followed in that strain by a set of pseudogenes (**Figure S1**). A BLASTX search showed that the two pseudogenes are probably derived from a protein-encoding gene mostly annotated as encoding a recombination-promoting nuclease or transposase. Interestingly, this gene was highly similar to SPV_2082 and an additional pseudogene, SPV_2340, located elsewhere on the D39V chromosome and also ComX-regulated. We speculate that the presence of a Repeat Unit of the Pneumococcus (RUP) (67) upstream of SPV_2082, in combination with the action of IS elements, might have enabled several duplication and/or reorganization events of the SPV_2082 locus. While these findings suggest that, in pneumococcal strains with an intact copy of this gene, it might be relevant to transformation and horizontal gene transfer, we do not expect *srf-01* (or pseudogenes SPV_2082 and SPV_2340, for that matter) to have a role in competence.

**Table 3.** ComX-regulated genes, distributed over 19 operons, as indicated by grey/white colored blocks. Secondary (under ‘Notes’) indicates either read-through after incomplete termination or the influence of an additional TSS. For complete information, including TSS positions, see **Table S5**. Purple and green cells indicate significance (*p* < 0.001, |^2^log FC| > 1; DESeq (45)). ^α^The *tyg* TSS was previously found to be ComX-regulated (28). An artificial 250 nucleotide transcript starting on this TSS was added to the annotation file to allow differential expression analysis. *Pseudogene.

| Gene | Product | TPM<br>0 min. | <sup>2</sup>log fold change |  |  | Notes |
| --- | --- | --- | --- | --- | --- | --- |
|  |  |  | 3 min. | 10 min. | 20 min. |  |
| *dut* | Deoxyuridine 5'-triphosphate nucleotidohydrolase | 78 | 2.7 | 1.5 | -0.2 |  |
| SPV_0028 | Hypothetical protein | 94 | 2.6 | 1.6 | -0.2 |  |
| *radA* | DNA repair protein | 96 | 2.5 | 2.4 | -0.1 |  |
| SPV_0030 | Carbonic anhydrase | 388 | 0.8 | 1.1 | 0.1 | Secondary |
| *srf-01* | ncRNA of unknown function | 71 | 3.7 | 3.3 | 0.9 | Putative pseudogene |
| SPV_0034* | IS1167 transposase | 3 | 3.6 | 3.7 | 1.1 |  |
| SPV_2082* | Hypothetical protein | 22 | 6.6 | 6.3 | 2.6 |  |
| *cibC* | CibAB immunity factor | 10 | 10.6 | 12.3 | 6.5 |  |
| *cibB* | Two-peptide bacteriocin peptide | 11 | 10.2 | 11.0 | 5.5 |  |
| *cibA* | Two-peptide bacteriocin peptide | 8 | 12.0 | 12.9 | 7.4 |  |
| SPV_2121 | Hypothetical protein | 124 | 4.6 | 4.4 | 0.6 |  |
| SPV_0186 | Competence-damage induced protein | 283 | 2.4 | 2.3 | 0.1 | Secondary |
| SPV_0683 | Hypothetical protein | 136 | 4.2 | 4.9 | 0.8 |  |
| *comEA* | Late competence DNA receptor | 4 | 9.8 | 9.2 | 3.7 |  |
| *comEC* | Late competence DNA transporter | 3 | 9.5 | 9.8 | 5.0 |  |
| SPV_2256 | Hypothetical protein | 156 | 4.0 | 4.8 | 1.4 | Secondary |
| SPV_2257* | ABC transporter ATP-binding protein | 91 | 3.7 | 5.0 | 1.3 | Secondary |
| SPV_0846 | Hypothetical protein | 68 | 3.9 | 5.4 | 1.5 | Secondary |
| *coiA* | Competence protein | 1 | 8.2 | 7.4 | 1.2 |  |
| *pepF1* | Oligoendopeptidase F | 252 | 1.3 | 1.2 | 0.0 | Secondary |
| SPV_0867 | O-methyltransferase family protein | 136 | 1.4 | 1.5 | 0.3 | Secondary |
| *radC* | DNA repair protein | 3 | 9.8 | 9.8 | 2.6 |  |
| *dprA* | DNA protecting protein | 6 | 10.1 | 8.8 | 4.6 |  |
| SPV_1308 | Oxidoreductase of aldo/keto reductase family, subgroup 1 | 182 | 0.9 | 1.4 | 0.3 | Secondary |
| *pgdA* | Peptidoglycan N-acetylglucosamine deacetylase | 397 | 1.4 | 1.7 | 0.1 | Secondary |
| SPV_2340* | Hypothetical protein | 222 | 2.5 | 2.6 | 0.2 | Secondary |
| *cclA* | Type IV prepilin peptidase | 2 | 9.3 | 9.0 | 3.7 |  |
| *ssbB* | Single-stranded DNA-binding protein | 9 | 10.7 | 11.9 | 6.8 |  |
| *lytA* | Autolysin/N-acetylmuramoyl-L-alanine amidase | 703 | 1.4 | 3.2 | 0.4 | Secondary |
| *dinF* | MATE efflux family protein | 288 | 2.2 | 3.8 | 0.4 | Secondary |
| *recA* | DNA recombination/repair protein | 391 | 3.0 | 3.8 | 0.7 | Secondary |
| *cinA* | ADP-ribose pyrophosphatase/nicotinamide-nucleotide amidase | 78 | 5.8 | 6.5 | 2.5 |  |
| *tyg*<sup>a</sup> | n/a | <1 | 6.7 | 6.8 | 4.2 |  |
| *yhaM* | 3'→5' exoribonuclease | 312 | 1.7 | 1.7 | -0.4 |  |
| *rmuC* | DNA recombination protein | 314 | 2.1 | 1.8 | -0.3 |  |
| SPV_1824 | ABC transporter permease protein | 34 | 0.9 | 2.7 | 0.2 | Secondary |
| SPV_1825* | IS630-Spn1 transposase | 162 | 0.5 | 1.7 | 0.0 | Secondary |
| *nadC* | Quinolinate phosphoribosyltransferase | 72 | 1.7 | 3.2 | 0.3 | Secondary |
| SPV_1828 | Hypothetical protein | 142 | 6.5 | 6.0 | 1.4 |  |
| SPV_2427* | S-adenosylmethionine-dependent methyltransferase | 2 | 10.3 | 13.0 | 7.2 |  |
| *comGG* | Late competence protein | 8 | 8.7 | 11.0 | 5.5 |  |
| *comGF* | Late competence protein | 2 | 9.9 | 12.2 | 6.5 |  |
| *comGE* | Late competence protein | 3 | 10.3 | 12.3 | 6.5 |  |
| *comGD* | Late competence protein | 4 | 10.1 | 12.1 | 6.3 |  |
| *comGC* | Late competence protein | 6 | 9.3 | 11.2 | 5.3 |  |
| *comGB* | Late competence protein | 8 | 9.4 | 10.8 | 5.4 |  |
| *comGA* | Late competence protein | 14 | 9.4 | 10.4 | 5.1 |  |
| *thiZ* | Thiamin ABC transporter ATPase component | 128 | 0.6 | 1.0 | -1.2 | Secondary |
| *thiY* | Thiamin ABC transporter substrate-binding component | 134 | 0.8 | 0.9 | -1.1 | Secondary |
| *thiX* | Thiamin ABC transporter transmembrane component | 88 | 0.8 | 1.2 | -1.1 | Secondary |
| SPV_2027 | Cytoplasmic thiamin-binding component of thiamin ABC transporter | 89 | 1.0 | 1.2 | -0.7 | Secondary |
| *cbpD* | Choline-binding protein D | 6 | 9.9 | 10.0 | 3.8 |  |
| *srf-29* | ncRNA of unknown function | 3 | 9.3 | 8.8 | 3.0 |  |
| *hpf* | Ribosome hibernation promotion factor | 624 | 0.9 | 1.4 | -0.3 | Secondary |
| *comFC* | Phosphoribosyltransferase domain protein | 3 | 8.6 | 8.5 | 2.7 |  |
| *comFA* | DNA transporter ATPase | 3 | 9.0 | 8.1 | 2.9 |  |

**Figure 6.**
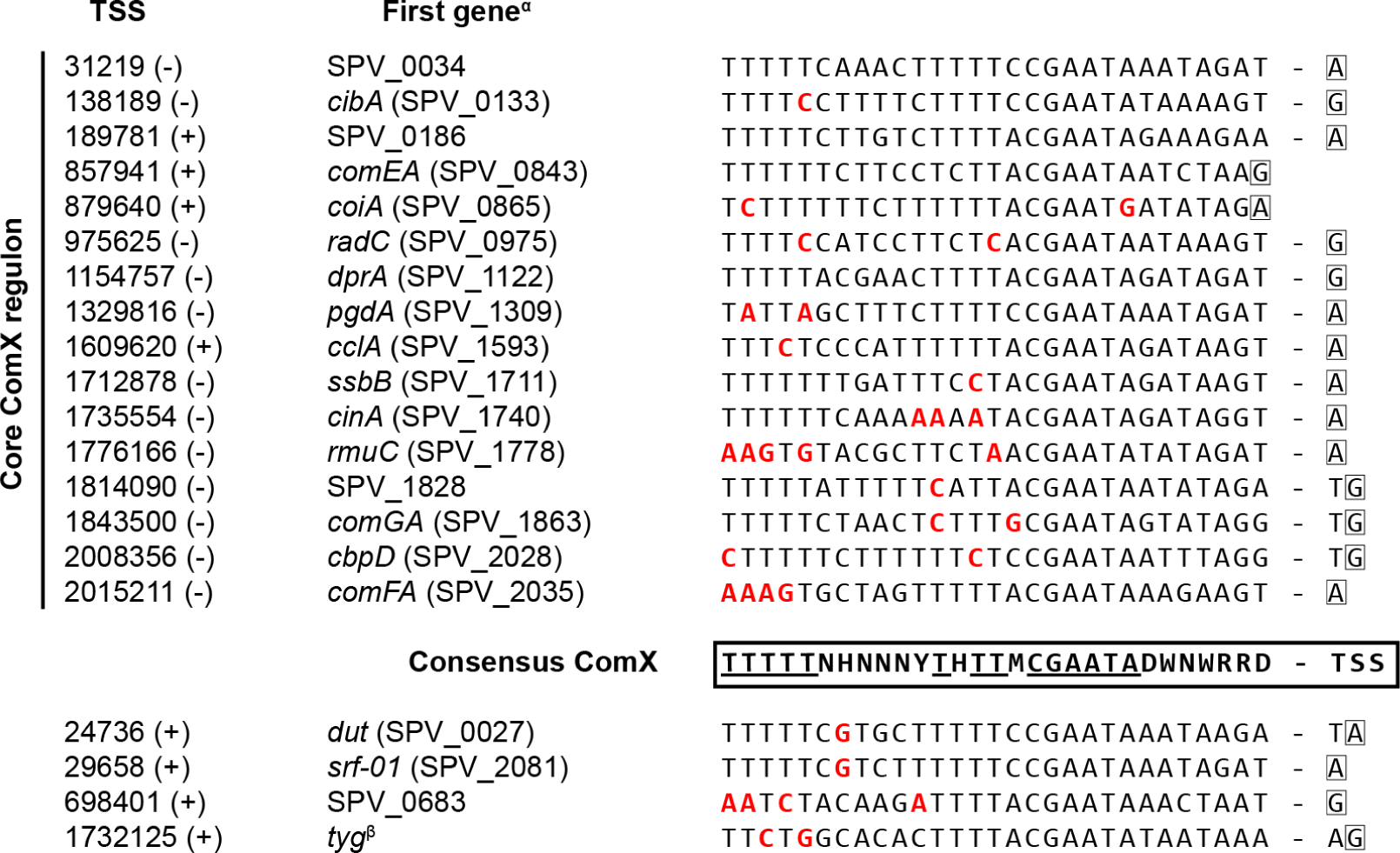
ComX-binding sequences on the *S. pneumoniae* D39V genome. Consensus sequence was determined as described in **Materials and Methods**. Multiple possible nucleotides are indicated, according to IUPAC nomenclature, by R (A/G), Y (C/T), W (A/T), M (A/C), D (A/G/T), H (A/C/T) or N (any). ^α^First gene both annotated in D39V and D39W (58), if available. ^β^*tyg* was described by Campbell et al. (28), its TSS was detected in PneumoBrowse (37), but no CDS or ncRNA has been reported (37, 66).

A second ncRNA, *srf-29*, is located upstream of and partially overlaps with *cbpD* (SPV_2028), a known late-*com* gene. It is not clear whether *srf-29* represents an uncharacterized RNA switch regulating *cbpD* expression, produces a functional sRNA, or simply is an artefact produced by a premature terminator (see PneumoBrowse coordinates 2008356-2008242 (-)).

Since TSS and terminator information permits a promoter-based interpretation of our data, we observed examples of complex operon structures, wherein TSSs or imperfect terminators inside the operon can lead to differences in expression between different genes in the same operon (35, 37, 68, 69). A striking example is the *cinA*-*recA*-*dinF*-*lytA* operon (SPV_1740-37), shown in **Figure 7**, which is under control of ComX, with only an inefficient terminator (27%) between *recA* and *dinF*. However, the presence of three internal TSSs, upstream of *recA, dinF* and *lytA*, respectively, leads to very different basal expression levels at t=0 (**Table 3**). Due to these differences, the effect size of competence induction on the expression of the four genes decreases from the 5’- to the 3’-end of the operon (**Figure 7**). Finally, Campbell et al. identified a transcription start site inside of and antisense to *dinF*, that was induced during competence and they provisionally named the associated hypothetical gene *tyg* (**Figure 7**; (28)). Although not discussed by Campbell and coworkers, both Håvarstein (70) and Claverys and Martin (66) argued that the peculiar positioning of the *tyg* TSS is reason for doubts regarding the functionality of any transcript produced. However, as Claverys and Martin concede, it cannot be ruled out that *tyg* has a role in mRNA stability of the *cinA*-*recA*-*dinF*-*lytA* operon. With Cappable-seq (52), we did indeed detect a transcription start site (37), accompanied by a consensus ComX recognition site (**Figure 7**). Since we did not detect a clearly demarcated associated transcript, we artificially annotated a 250 nucleotide long transcript, starting at the *tyg* TSS, to allow differential expression analysis. The time-dependent expression trend of this transcript during competence seemed to follow that of other late-*com* genes (35). However, the extremely low detected expression level prior to and, even, after CSP addition precluded any further statistical analysis regarding differential expression or clustering.

**Figure 7.**
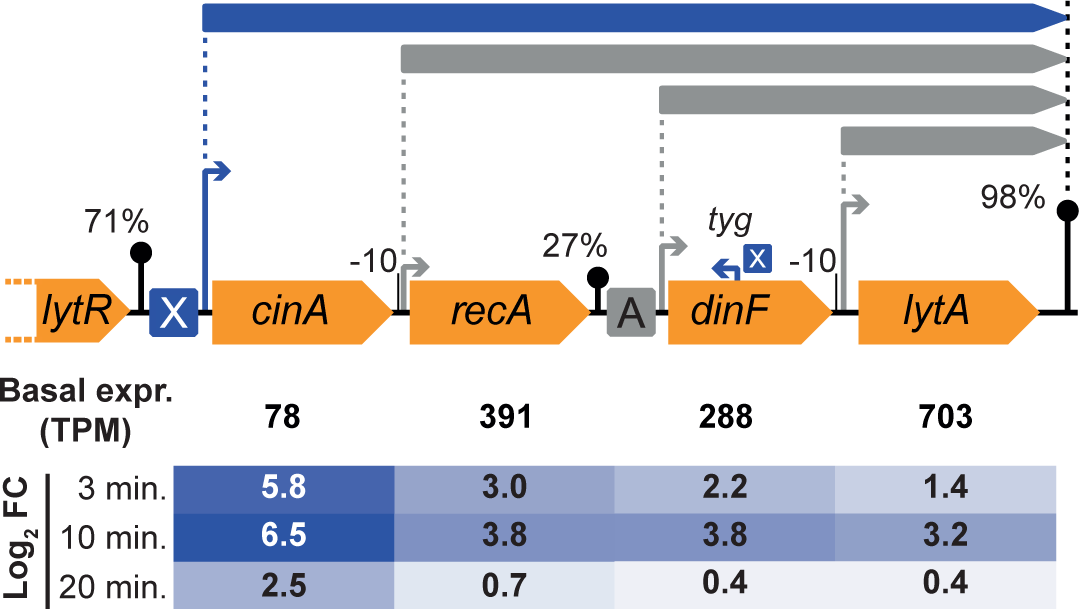
Top: overview of the complex *cinA*-*recA*-*dinF*-*lytA* operon, with an imperfect internal terminator and TSSs upstream of each gene, leading to four overlapping operons. The TSS upstream of *cinA* is preceded by a ComX-binding site and addition of CSP indeed affects expression of all four genes in the operon. Bottom: ^2^log(fold change) relative to t=0 (i.e. basal expression). However, the effect size decreases with every gene, due to differences in basal expression from the internal TSSs. Additionally, a ComX-regulated TSS, is found inside of and antisense to *dinF*, giving rise to the hypothetical transcript *tyg* (28), with an unknown 3’-end.

The expression pattern of 44 out of the 56 members of the ComX-associated WGCNA cluster (cl. 11) can now be linked to a ComX-regulated TSS. Bearing in mind that the clustering was performed based on expression throughout all 22 infection-relevant conditions studied in PneumoExpress (35), only five other cluster members (SPV_0553, SPV_0957-59 and SPV_2317) showed an expression pattern similar to ComX-regulated genes in competence conditions specifically. While the TSS for SPV_0553 has not been determined, this gene is surrounded by two Repeat Units of the Pneumococcus (67) and one BOX element (60) and nothing resembling a ComX-binding site was found near it. Secondly, SPV_2317 represents a novel ncRNA (*srf*-*19*), potentially an RNA switch, that is preceded by a predicted RpoD site, rather than a ComX site. The last cluster member, operon SPV_0957-59, contains *rpoD* (SPV_0958). In light of the proposed role of RpoD in the shutdown of late competence (see above), it would be interesting if its upregulation was directly induced by ComX. However, analysis of the promoter region of the operon yielded no indication of a ComX-binding site and the mechanism of *rpoD* induction in competence continues to elude us.

### The CiaR regulon is induced during competence and contains a novel non-coding RNA

Besides early and late-*com* genes, terminology reserved for the ComE- and ComX regulons, respectively, many other genes are more indirectly affected by the addition of CSP. Although a small portion of these genes can already be seen to be affected after 3 minutes, we will refer to all of these genes collectively as ‘delayed’, as these changes occur at least after the activation of the ComE regulon and most likely also downstream of the ComX regulon. Among the delayed genes are nearly all members of the CiaR regulon (**Table 4**) and promoter analysis of the core members of the CiaR regulon (**Table S3**, **Figure S2**) returned the CiaR recognition site as previously reported (30, 48, 49): [TTTAAG]-[N_5_]-[TTTAAG]-[N_22_]-[TSS]; **Figure 3B**, right). Analysis of other affected promoters turned up three additional monocistronic operons (*ccnC*, SPV_0098 and SPV_0775), all of which have already previously been reported to be CiaR-regulated. While SPV_0098 is expressed from two different TSSs (48) and *ccnC* (SPV_2078) expression is affected by transcriptional read-through from the upstream ComE-regulated *comW* operon, it is not clear why SPV_0775 does not cluster with other CiaR-regulated genes.

**Table 4.**
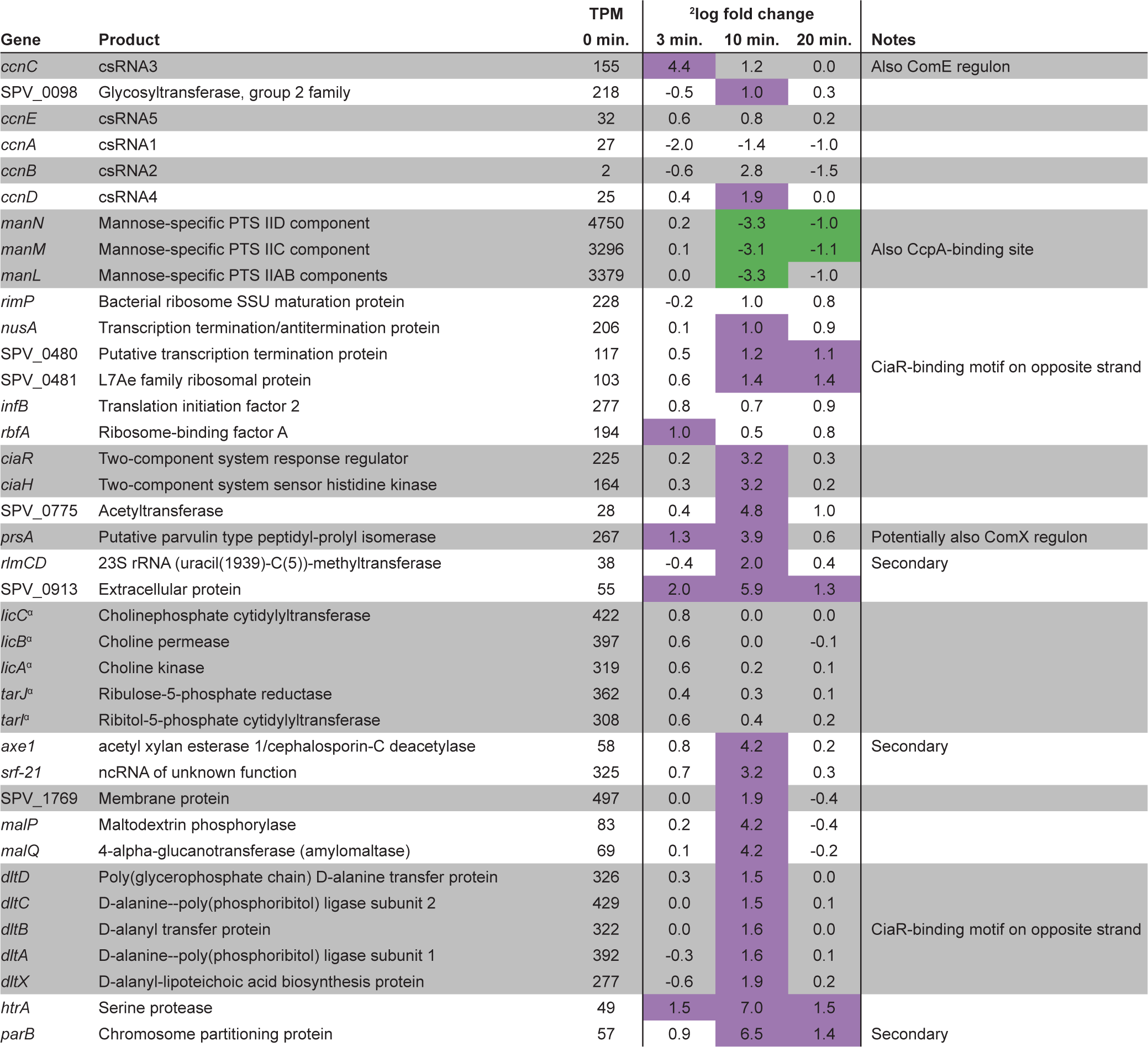
CiaR-regulated genes, distributed over 18 operons, as indicated by grey/white colored blocks. Secondary (under ‘Notes’) indicates either read-through after incomplete termination or the influence of an additional TSS. For complete information, including TSS position, see **Table S5**. Purple and green cells indicate significance (*p* < 0.001, |^2^log FC| > 1; DESeq (45)). ^α^Operon has two different detected TSSs. The TSS at 1159217 (-) is under control of CiaR.

Since CiaR-binding sites were found on the opposite strand for *dltXABCD* (SPV_2006-02; upregulated) and *manLMN* (downregulated), we speculate that operon SPV_0478-83 (i.a. *rimP, infB, nusA* and *rbfA*), encoding several proteins involved in translation, is also regulated by CiaR (**Figure S2**). Intriguingly, another new member of the CiaR regulon is *srf-21* (SPV_2378), a novel, uncharacterized non-coding RNA (37). The TSS from which this ncRNA is expressed was already part of the reported CiaR regulon, but was linked to the overexpression of the downstream *axe1* gene (SPV_1506). Inspection of the transcriptional layout of the region (**Figure 8A**) shows that *srf-21* and *axe1* are separated by a relatively efficient terminator and a second TSS. Nonetheless, *axe1* overexpression might still be attributed to read-through from *srf-21*. We did not find any similar ncRNAs in RFAM and BSRD databases (71, 72) and, since *axe1* is expressed from its own TSS, it seems unlikely that *srf-21* functions as an RNA switch. Preliminary minimum free energy (MFE) secondary structure prediction with RNAfold (73) and target prediction with TargetRNA2 (74) did not provide us with any clear hints with regard to the function of this ncRNA. Firstly, the predicted MFE structure (**Figure 8B**) might only represent a transient conformation, since it makes up less than 1% of the modelled ensemble. Secondly, sRNA target prediction produced many potential targets (**Table S4**). Some candidate regions are less likely to be targeted, because they are located more than 20 nucleotides upstream of the start codon or even upstream of the TSS, ruling out a possible interaction between *srf-21* and the transcript in question. However, future work will be necessary to reveal whether any of the remaining genes (e.g. *queT, pezA* or *cps2H*) are regulated by *srf-21* or, indeed, whether this ncRNA might have a completely different mode of action.

**Figure 8.**
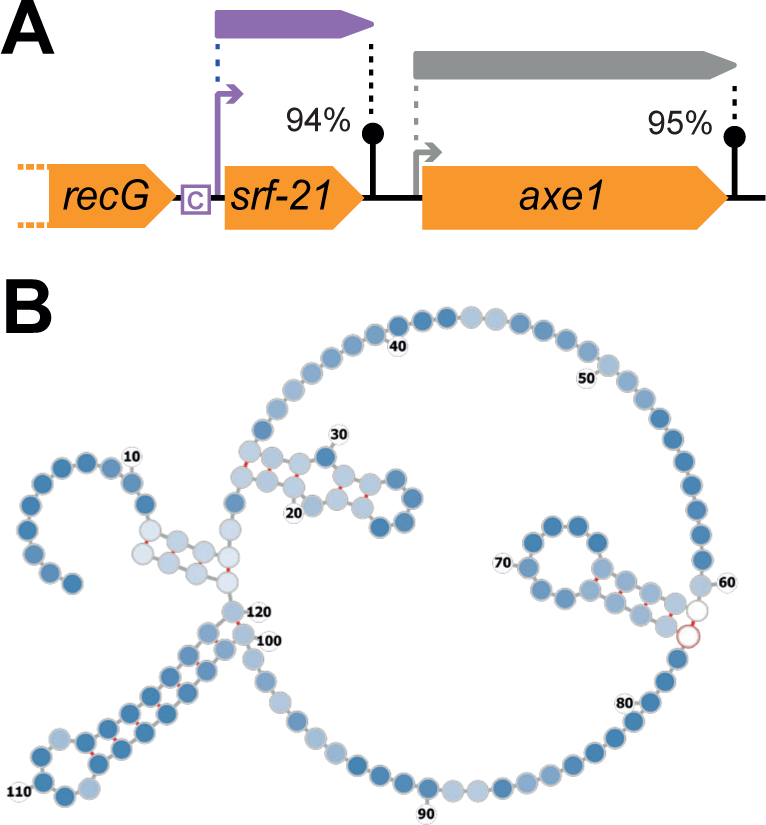
Non-coding RNA *srf-21* (SPV_2378) is part of the CiaR-regulon. (**A**) Overview of the genomic context of *srf-21*. CiaR-dependent upregulation of downstream gene *axe1* might be due to read-through from *srf-21*. The CiaR-binding sequence is indicated by a boxed ‘C’. (**B)** MFE secondary structure of *srf-21*, as predicted by RNAfold (73).

It is noteworthy that four reported promoters of the CiaR regulon were not found to be significantly affected during competence. Firstly, the *tarIJ-licABC* operon is under the control of two TSSs and thereby apparently less sensitive to CiaR control. Finally, three out of five csRNAs, described by Halfmann et al. (30), did not appear to be significantly affected (**Table 4**). It should be noted, however, that the statistics on these short transcripts is rather poor and the current data can neither support nor contradict their upregulation in competence. However, we do believe that the data presented by Halfmann et al. regarding CiaR-regulation seem perfectly convincing, and the promoter regions of each of the five csRNAs contain a clear match with the consensus CiaR-binding site (**Figure S2**).

### Other known regulons affected during competence

Still unknown in previous descriptions of the competence regulon, the VraR (LiaR) regulon has been described by Eldholm et al. to be activated in response to competence-induced cell wall damage (15). Based on three pneumococcal promoters and six *L. lactis* promoters (**Table S3**, **Figure S3**), we rebuilt the consensus motif described by Eldholm et al. (**Figure 4**) and observed that all of these motifs are located 32-34 nucleotides upstream of the corresponding TSS. In contrast, we could confirm the reported presence of a VraR-binding site upstream of *hrcA* (15), but this site is 81 nucleotides removed from its target TSS. However, the fact that this promoter region also carries two HrcA-binding sequences could account for this difference in spacing. Finally, we suggest that SPV_1057 (spr1080 in R6) and SPV_1160 (spr1183) are not regulated by VraR, contrary to the report by Eldholm and coworkers. Firstly, both of these genes lacked a detected TSS and neither was found to be differentially expressed in our study. Secondly, the reported recognition site for SPV_1057 is actually located downstream of the gene, inside a repeat region (ISSpn7 element). Finally, as recognized by Eldholm et al., SPV_1160 represents a 5’-truncated version of a gene putatively encoding the ATP-binding component of an ABC transporter. These observations, combined with the fact that the reported VraR-binding site is located only 24 nucleotides upstream of the annotated start of SPV_1160, led us to conclude that SPV_1160, like SPV_1057, is not regulated by VraR. This limits the VraR regulon to 15 genes, distributed over 4 operons (**Table S5**).

Regardless of whether or not competence should be regarded as a stress response mechanism in itself, it is clear that the activation of competence, at least indirectly, leads to a multifactorial stress response. Besides VraR and CiaR, also the well-characterized HrcA (3, 75) and CtsR (3, 76) regulons are activated in competent cells, as previously reported (3). The only addition to be made here regarding the HrcA regulon is the annotation of a gene encoding a protein of unknown function (SPV_2171), not previously annotated in D39 or R6 strains. This gene is located between *dnaK* (SPV_0460) and *dnaJ* (SPV_0461) and therefore regulated by both VraR and HrcA (**Table S5**). Also the CtsR regulon was found to be upregulated almost entirely. Only for *clpP* (SPV_0650), the observed upregulation was below the employed cutoff, possibly because its basal expression level (t=0) is 4- to 40-fold higher than that of other *clp* genes. Finally, the upregulation of an uncharacterized two-component regulatory system (SPV_2020-19), with unknown consequences, could be attributed to transcriptional read-through from the upstream *ctsR*-*clpC* operon.

Two other regulons seemed overrepresented in the set of differentially expressed genes (*p* < 10^-4^, hypergeometric test). Firstly, all six genes predicted to be regulated by GntR (SPV_1524), as based on homology to *Streptococcus pyogenes* Spy_1285 (54), were found to be upregulated 10 and 20 minutes after addition of CSP. These six genes are distributed over two operons (SPV_0686-88 and SPV_1524-26). Secondly, a significant number (31) of RpoD-regulated genes were downregulated, mostly after 10 and 20 minutes, which may readily be explained by the competition for RNA polymerase of RpoD (σ^A^) with the alternative sigma factor ComX (σ^X^).

### Other differentially expressed genes

A total of 367 genes (i.e. 17% of all annotated genes) are either found to be differentially expressed or, at least, to be under the control of a TSS that appears to be differentially regulated at some point during competence (**Table S5**). The response of a large portion (204 genes) of these can be ascribed to the action of one of the transcriptional regulators discussed above. While 56% of the latter group display a maximum absolute change in expression of more than fourfold, only 29 of the remaining 163 genes (16%), distributed over 14 operons, meet the same criterion. These data show that the bulk of strong induction or repression can be explained by a small set of regulators. Among the 29 strongly differentially expressed genes with no known regulators, our data confirmed the upregulation of *rpoD*, in line with the role that RpoD might play in late competence shutdown (33).

Functional analysis did not reveal any Gene Ontology or KEGG classes overrepresented among upregulated genes. Only classes related to ribosomes and translation seemed overrepresented, but the realization that all affected genes from these classes were part of a single operon (shown in **Table 1**) led us to discard them due to lack of evidence (see **Materials and Methods**). Similarly, most potential hits among downregulated genes were discarded. Only classes related to thiamine metabolism (GO:0009228, KEGG:ko00730) remained. The five genes in question are distributed over four operons: *adk* (SPV_0214), *thiM-1-thiE-1* (SPV_0623-24), *thiD* (SPV_0632), and *sufS* (SPV_0764). Two of these four operons are regulated by a TPP riboswitch, an RNA element that, when bound to thiamine pyrophosphate (TPP), prevents transcription of the downstream operon (77). We suspect, therefore, that the temporary growth lag accompanying competence development (3) leads to a transient accumulation of TPP, which then represses transcription of operons under control of a TPP riboswitch. Indeed, both other D39V operons led by a TPP riboswitch show a similar expression trend 10-20 minutes after competence induction. Firstly, *ykoEDC-tenA-thiW-thiM-2-thiE-2* (SPV_0625-31) is already very lowly expressed prior to CSP addition, preventing significant downregulation (not shown). The second operon, SPV_2027-*thiXYZ* (SPV_2027-24), was excluded from gene enrichment analysis since it was part of the ComX regulon (**Table 3**). Since the hypothesized accumulation of TPP seems to occur with a delay, relative to the activation of the ComX regulon, these genes are first upregulated (3-10 min) and then downregulated (20 min), even relative to the basal expression level.

### Comparison to previous reports of the competence regulon

Finally, we compared our findings with previous, microarray-based studies by Peterson et al. (4) and Dagkessamanskaia et al. (3). Although both studies give a remarkably complete overview, our approach allowed us to refine and nuance the description of the competence regulon even further (**Table S5**). The higher sensitivity and accuracy of Illumina sequencing, the improved genome annotation and the application of a promoter-based analysis (rather than gene-based) allowed us to expand the set of genes under direct control of ComE and ComX to 40 and 55 genes, respectively (combined: 4% of all genes). Especially several genes with putative BlpR-binding sites (**Table 2**), and therefore a generally weaker response, were missing from previous reports. Additionally, the previously reported *briC* operon (36) is now included in the ComE regulon and we confirmed that, while undetected by Dagkessamanskaia et al., *ybbK* and *def2* (early) and *radC* (late) are indeed part of the *com* regulon (**Tables 2** and **3**). Remaining discrepancies could be explained either by transcriptional read-through or the absence of certain elements (e.g. ncRNAs) from the TIGR4 and R6 genome annotation files used by Peterson et al. and Dagkessamanskaia et al., respectively.

Not surprisingly, larger discrepancies were found for genes displaying delayed differential expression, since their fold changes (mostly) are considerably smaller and therefore more sensitive to technical variation and differences in experimental conditions (e.g. growth medium, pH or time of induction). In contrast with the previous studies, we recovered nearly the entire CiaR regulon as differentially expressed in our data, confirming the high sensitivity of RNA-seq, compared to microarray-based technology.

## DISCUSSION

Competence for genetic transformation is defined as a state in which a bacterial cell can take up exogenous DNA and incorporate it into its own genome, either in the form of a plasmid or via homologous recombination. Since the very first demonstrations of bacterial transformation were provided by Griffith and Avery et al. (78, 79) in *S. pneumoniae*, the pneumococcal competence system has been widely studied as soon as the required tools became available. Therefore, much knowledge has been assembled about how this state is regulated, which environmental triggers affect its development and what downstream consequences it has. Indeed, several studies have been performed to compile a comprehensive list of all competence-regulated pneumococcal genes. The most recent of these studies (3, 4), although of very high quality and invaluable to the research field, date from nearly fifteen years ago. Since then, the fields of transcriptome analysis and genome sequencing and annotation have been revolutionized by second-(e.g. Illumina) and third-generation (e.g. PacBio) sequencing techniques. Therefore, we have analyzed Illumina-based RNA-seq data (35), using the recently sequenced and deep-annotated *S. pneumoniae* D39V strain (37), to refine the previously reported pneumococcal competence regulon. Additionally, rather than just reporting affected individual genes, we used previously determined transcript boundaries (TSSs and terminators; (37)) to identify the affected promoters, which may be used to gain more insights into the regulatory processes at work during competence.

In short, we report that ComE directly regulates 15 early-*com* transcripts (40 genes), including 4 transcripts (10 genes) that are, probably, part of the BlpR regulon (**Table 2**, **Figure 5**). Alternative sigma factor ComX (σ^X^) was found to control 19 late-*com* transcripts (55 genes), in addition to the previously described *tyg* TSS, inside and antisense to *dinF* (**Table 3**). We should note that four genes from the early and late-*com* regulons (e.g. *blpC*) did not meet fold-change and/or statistical cutoff values, but were part of operons that were clearly regulated. For each of these genes, the observed expression trends correlated with those of other operon members.

Our data confirmed that, as shown in the previous studies (3, 4), the activation of the early and late-competence regulons indirectly resulted in the activation of several other regulons, most of which are implicated in pneumococcal stress response. Firstly, out of the newly compiled CiaR regulon (18 transcripts, 38 genes), based on the work of Halfmann et al. (30), 4 operons were not found to be significantly affected (**Table 4**). Other affected regulons were those under control of VraR (LiaR), HrcA, CtsR and GntR (**Table S5**). Additionally, genes led by a TPP riboswitch, were downregulated, suggesting a transient increase in intracellular thiamine pyrophosphate levels in competent cells.

Many other genes, including *rpoD*, are up-or downregulated through unknown mechanisms. In total, approximately 140 transcripts (containing 367 genes, 17% of all genes) undergo some extent of differential expression. For several reasons (e.g. fold changes near cutoff, expression from multiple TSSs or poor statistics due to low expression), 79 genes from these operons did not individually meet the detection criteria, leaving 288 differentially expressed genes (13%) when following the traditional, gene-based analysis approach (**Figure 2**). Among the affected genes are several small, non-coding transcripts. Some of these ncRNAs, like the CiaR-activated csRNAs (see below), have been characterized and we showed that *srf-01* is unlikely to be functional. For others, e.g. *srf-03* (upstream of *comAB*) and *srf-21* (upstream of *axe1*), future work is required to determine their role, if any, during competence.

Given the fact that so many different functionalities are activated during competence, including stress response systems such as the Clp protease and several chaperone proteins, Claverys et al. proposed to refer to the system more neutrally as ‘X-state’ (for ComX). We would argue, however, that the primary response to a high extracellular CSP level is the activation of the ComE and ComX regulons, which mostly encode proteins relevant to transformation. Firstly, the expression of fratricin CbpD (SPV_2028), along with immunity protein ComM (SPV_1744), allows for the lysis of neighboring non-competent cells, which may offer access to both nutrients and DNA (80, 81). The upregulation of the gene encoding autolysin LytA (*lytA*) would fit in nicely here, since the simultaneous deletion of *lytC* and *lytA* abolishes competence-induced lysis completely (80). However, basal level *lytA* expression was reported to be already sufficient for the observed lysis rate (82). Next, the DNA uptake machinery comes into play, involving many proteins encoded by the late-*com* genes, as reviewed by Claverys et al. (16); most of ComGC-ComGG (SPV_1861-57), ComEA/C (SPV_0843-44) and ComFA (SPV_2035) are, together with constitutively expressed endonuclease EndA (SPV_1762) and through the action of prepilin peptidase CclA (SPV_1593) and proteins ComGA/B (SPV_1863-62), assembled into a pilus-like structure (83). The import of DNA is followed by DNA processing and recombination, involving DNA protection protein DprA (SPV_1122), recombinase RecA (SPV_1739), ssDNA-binding SsbB (SPV_1711) and other late-competence proteins (84–87). For the reasons discussed above, combined with the fact that the vast majority of other differentially expressed genes display a more modest change in expression (**Table S5**), we decided to keep calling the system competence for genetic transformation. We do, however, agree that the downstream effects of competence development should not be ignored, as discussed below.

The genome of *Streptococcus pneumoniae* D39V contains 13 two-component regulatory systems (TCSs) (37, 88). Interestingly, the regulons of two TCSs (ComDE and BlpRH) are activated during competence, while the regulons of another two TCSs (CiaRH and VraRS) are activated shortly after. A fifth, uncharacterized TCS (SPV_2020-19) is also slightly upregulated, possibly due to transcriptional read-through from the *ctsR-clpC* operon. The additional activation of stress-related regulons of HrcA and CtsR grant support to the hypothesis, as proposed by Prudhomme at al. (5), that competence activation serves as a general stress response in the pneumococcus, which lacks the SOS response that is common is many other bacteria. The activation of competence in response to various types of stress (5–8) provides even more relevance to this idea. On the other hand, Dagkessamanskaia et al. showed that a deletion of the CiaR regulon causes an extended growth lag, as well as a stronger activation of the HrcA and CtsR regulons, after competence induction (3). Both observations are in line with the notion that the development of competence is accompanied by a significant burden to the cell. It seems likely, therefore, that the main benefit of the activation of the CiaR and other stress response regulons during competence is to deal with the stress invoked by competence itself, by building the large membrane-and cell-wall-protruding DNA-uptake machinery and the production of cell wall hydrolases such as the fratricin CbpD. It is not unthinkable, however, that the activation of the many different stress response regulons renders it beneficial to a pneumococcal cell to become competent even in specific stressful conditions that do not require DNA uptake or recombination machineries. Whether or not such conditions played a role in the (co-)evolution of competence and downstream processes is open to speculation. In this respect, it is interesting to note that the production of competence-induced bacteriocins was found to be important to prevent ‘intruder’ pneumococci from colonizing (89).

The apparent severity of the stress imposed on a competent cell emphasizes the need to shut down competence after a short transformation-permissive time window. In addition to the known role of DprA in early competence shut-down, Weyder et al. proposed that the upregulation of *rpoD* is responsible for the, less-efficient, shut-down of late competence (33). Related to this, RpoD-regulated genes are, to some extent, overrepresented among downregulated genes during competence. While this explains the need for upregulation of *rpoD*, the underlying mechanism is still unknown. Other aspects that might play a role in the shutting down of competence are the CiaR-mediated upregulation of HtrA, which has been shown to degrade extracellular CSP (9), and the recently discovered CiaR-regulated non-coding csRNAs (*ccnA-E*) (30, 90, 91), which were shown to repress ComC translation in an additive fashion.

Finally, although several of the activated regulons are quite well-understood, still a large portion of affected genes are differentially expressed through unknown mechanisms. It seems plausible that many of these are due to the sudden and severe shift in metabolic state. For example, the higher translational demands during competence could lead to the upregulation of genes encoding ribosomal proteins (**Table 1**). Similarly, the hypothetical transient increase in TPP concentrations, leading to riboswitch-mediated downregulation of four operons, could be accompanied by the accumulation or depletion of other, unknown metabolites, with potential transcriptional consequences.

## Supporting information

## DATA AVAILABILITY

The data analyzed here can be extracted from PneumoExpress (https://veeninglab.com/pneumoexpress-app). Raw RNA-seq data used to build PneumoExpress was deposited to the GEO repository: accession number GSE108031.

## ACKNOWLEDGMENTS

We are grateful to A. de Jong and S. Holsappel for (bio)informatics support; and S.B. van der Meulen for sharing *L. lactis* TSS data.

