## Supplementary material for "Refining the pneumococcal competence regulon by RNA-sequencing"

### Supplementary table legends

**Table S1**. Expression values (TPM) of all *S. pneumoniae* D39V genes.

**Table S2**. Expression clusters, as determined by weighted gene co-expression network analysis (WGCNA; (1)).

**Table S3**. Transcription start sites used to create position weight matrices for recognition sites of ComE, ComX, CiaR and VraR.

**Table S4**. Genes putatively targeted by *srf-21*, as predicted by TargetRNA2 (2). Target coordinates have been modified to match D39V, rather than old D39W annotation (3). Where possible, distance relative to the transcriptional start site is given. Interaction sites further than 20 bps upstream of the start codon or largely upstream of the TSS are shown in blue and red, respectively. The candidate region upstream of *queT* (SPV_0898; orange) is inside the reported preQ1-II leader (4).

**Table S5**. All differentially expressed genes in response to CSP addition. Reports in literature were used to predict putative regulation by HrcA (5), CtsR (6), GntR (From RegPrecise; (7)) and/or a PyrR RNA switch (8, 9). Purple and green cells indicate significance (*p* < 0.001, |^2^log FC| > 1; DESeq (10)). ^α^Expression of *ccnC* is controlled by both ComE (TSS at 22142 (+)) and CiaR (TSS at 23967 (+)). ^β^Operon has two detected TSSs. The TSS at 183663 (+) is under control of VraR. ^γ^Expression of SPV_2027-*thiXYZ* is controlled both by ComX (TSS at 2008356 (-)) and a TPP riboswitch (TSS at 2006885 (‑)).

**Table S6**. Gene Ontology and KEGG classifications and predicted regulons used for gene enrichment analysis, as reported in PneumoBrowse (8).

### Supplementary figures

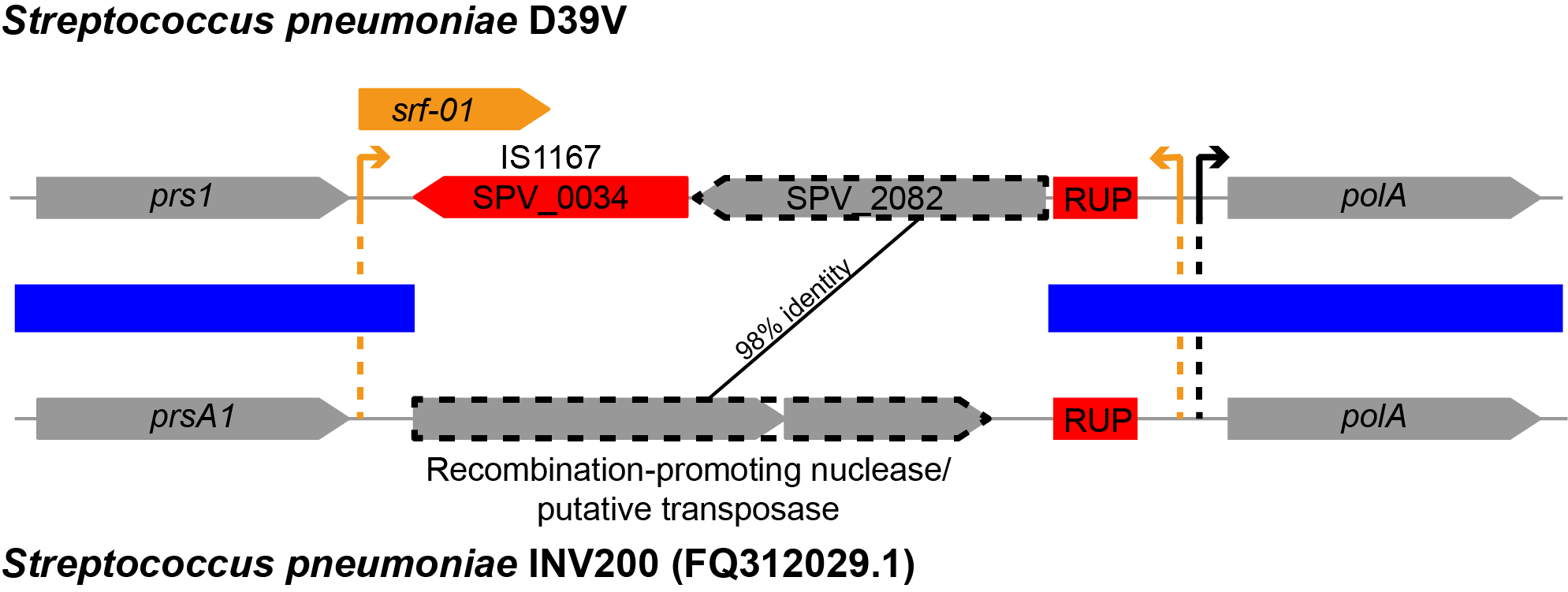

**Figure S1**. Comparison of *S. pneumoniae* D39V *srf-01* (SPV_2081) region and the corresponding region on the *S. pneumoniae* INV200 genome (GenBank: FQ312029.1). Blue blocks indicate conserved sequences, orange arrows indicate ComX-regulated transcription start sites, while the black arrow indicates an RpoD-regulated transcription start site. Red blocks represent repeat-containing sequences.

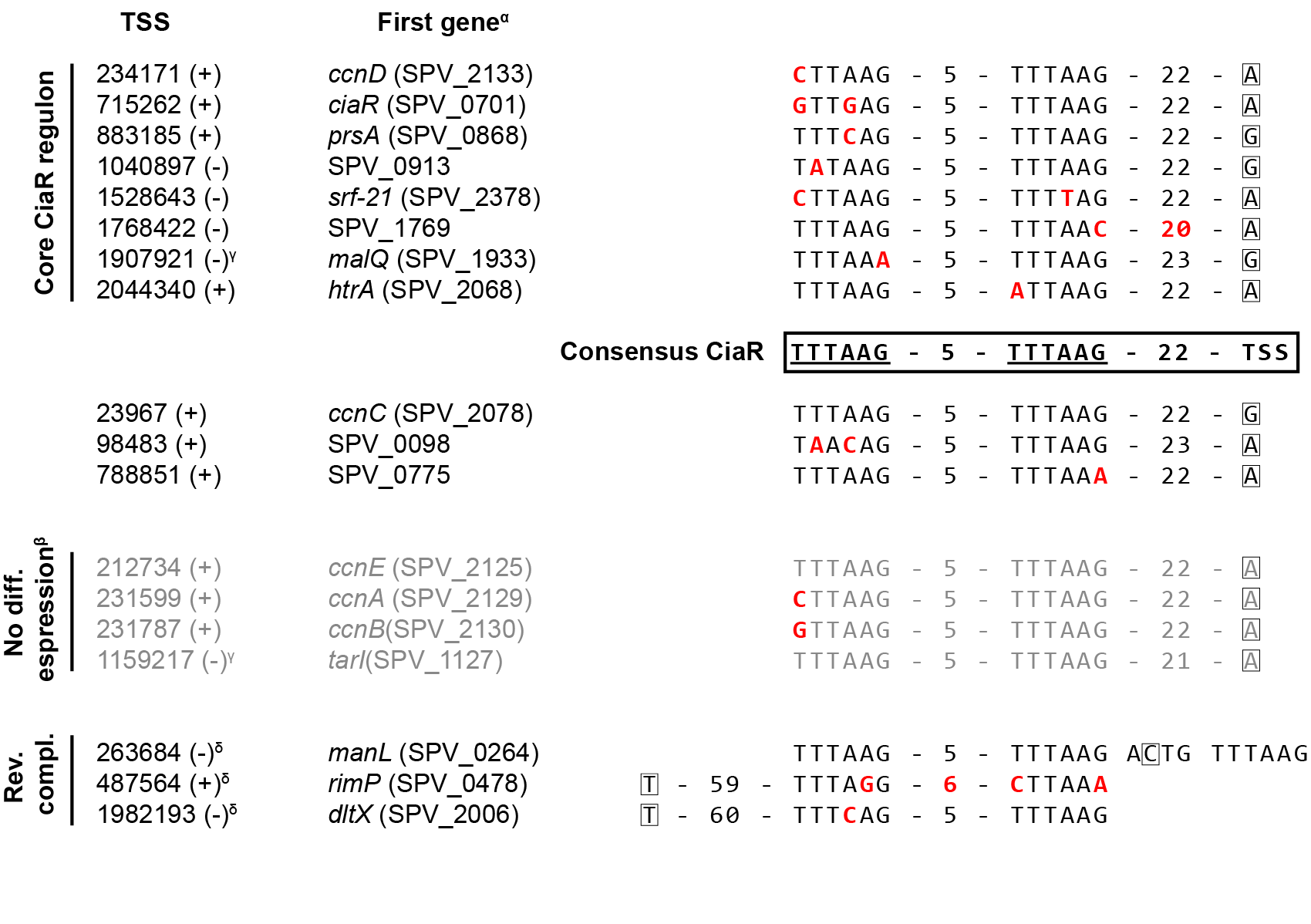

**Figure S2**. CiaR-binding sequences on the *S. pneumoniae* D39V genome. Consensus sequence was determined as described in **Materials and Methods**. ^α^First gene both annotated in D39V and D39W (3), if available. ^β^See **Table 4**. ^γ^The *tarIJ-licABC* and *malQP* operons are each under control of two different TSSs, one of which is regulated by CiaR. ^δ^Displayed sequences are reverse complementary to the transcription direction. In the case of *manLMN*, this leads to CiaR-mediated repression rather than activation.

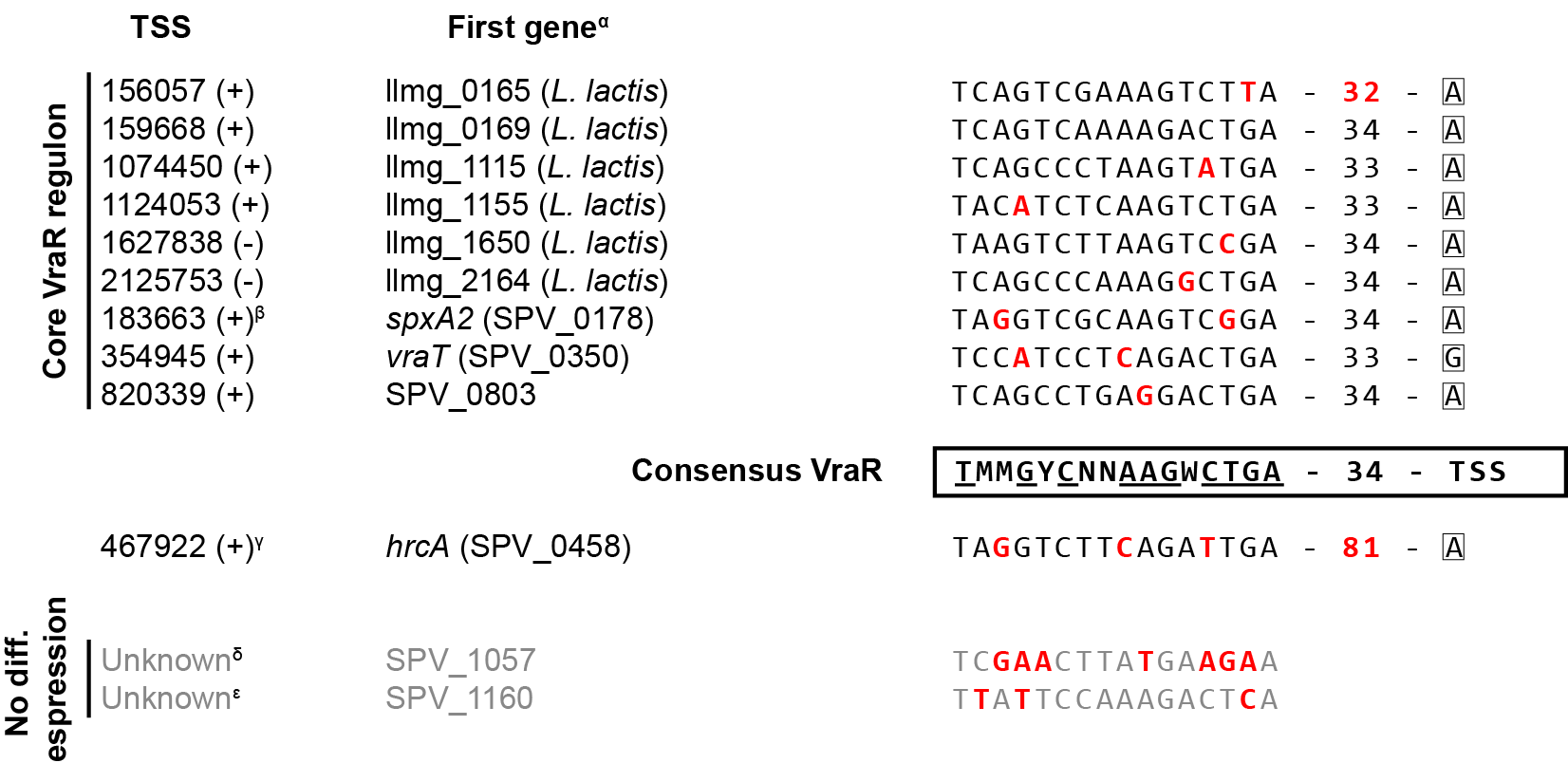

**Figure S3**. VraR-binding sequences on the *S. pneumoniae* D39V and *L. lactis* genome. Consensus sequence was determined as described in **Materials and Methods**. Multiple possible nucleotides are indicated, according to IUPAC nomenclature, by Y (C/T), W (A/T), M (A/C) or N (any). ^α^First gene in *L. lactis* or first gene both annotated in D39V and D39W (3), if available. ^β^Gene *spxA2* is under control of two different TSSs, one of which is regulated by VraR. ^γ^The *hrcA* operon is coregulated by HrcA and VraR, possibly accounting for the different spacing to the TSS. ^δ^Reported site (11) is located downstream of SPV_1057, inside a transposon sequence. ^ε^Distance between reported site (11) and annotated start codon of SPV_1160 is 24 nucleotides. Moreover, SPV_1160 is a 5’-truncated pseudogene.
